# Contribution of statistical learning to learning from reward feedback

**DOI:** 10.1101/2024.04.27.591445

**Authors:** Aryan Yazdanpanah, Michael Chong Wang, Ethan Trepka, Alireza Soltani

## Abstract

Natural environments are rich with patterns and regularities. Thus, it is not surprising that most animals have evolved neural mechanisms to detect and adapt to these regularities. Such detections and adaptations can, in turn, influence several cognitive functions including attention, perception, and memory, thereby enhancing survival. Here, we investigated whether detecting environmental regularities that are irrelevant to obtaining rewards can influence learning about multi-dimensional choice options––a process often constrained by the scarcity of reward feedback. To explore this, we trained human participants to perform a multidimensional reward-learning task alongside an orthogonal sequence-prediction task. We found that although feature-specific regularities in the sequence-prediction task were not predictive of reward, they incidentally biased participants’ behavior toward the feature with regularity during the reward-learning task. Fitting choice behavior with various computational models revealed that this effect was more consistent with modulations in learning rather than decision making, as evidenced by higher learning rates for this feature. This was particularly apparent for learning from chosen, rewarded options and unchosen, unrewarded options, demonstrating that environmental regularities can amplify confirmation bias in reward learning. Our results thus extend the notion of confirmation bias in learning about options and actions to their features. Furthermore, they suggest that temporal regularities can influence reward learning by biasing the association of reward with specific features and by enhancing confirmation bias. These effects help reduce dimensionality of the learning task to mitigate the curse of dimensionality in reward learning.

## Introduction

While constantly changing, natural environments exhibit numerous spatial and temporal regularities across multiple timescales (Soltani et al., 2021). For instance, tree fruits change color from green to yellow, orange, and red during the summer and fall, while their size varies significantly. Detecting and learning environmental regularities is ecologically crucial because it enhances predictability of the environment and reduces its complexity, allowing animals to navigate and adjust their behavior more efficiently (Gibson, 1969). Not surprisingly, our brain is adept at identifying relationships between events, cues, and stimuli, a process termed statistical learning (Saffran et al., 1996; Turk-Browne et al., 2009). This ability has been shown to enhance many cognitive functions including object categorization (Turk-Browne et al., 2010), short-term memory (Umemoto et al., 2010), auditory perception (Saffran et al., 1999), language acquisition, and social cognition (Conway & Christiansen, 2001; Saffran & Kirkham, 2018).

Ultimately, however, survival hinges on the ability to learn and select options and actions that yields reward, commonly referred to as reward learning. In naturalistic environments, however, reward learning can be very challenging because numerous choice options and/or actions precede a single reward feedback, while each option possesses multiple features or attributes. For example, fruits have many features, such as color, texture, and size. This complexity causes the required sample size for effective learning to grow exponentially, leading to the so-called curse of dimensionality (Sutton & Barto, 2018). One possible solution to mitigate the curse of dimensionality is to learn the predictive values of features to estimate reward values associated with individual objects or choice options (Farashahi et al., 2017; Leong et al., 2017; Niv et al., 2015). Such feature-based learning provides a good candidate for learning reward contingencies in multidimensional reward environments as it can reduce dimensionality without compromising too much precision (Farashahi et al., 2017, 2018, 2020). But how are these features identified among many features in the first place?

Notably, unlike reward learning that requires specific events (e.g., reward feedback), statistical learning can happen continuously as we receive the stream of sensory stimuli from our environment. Indeed, humans can automatically extract multiple statistical features of regularities without explicit instruction (Fiser & Aslin, 2002), and statistical learning can function as a form of associative learning similar to that required for reward learning (Turk-Browne et al., 2009). These raise the question of whether statistical learning could enhance reward learning by mitigating the curse of dimensionality.

A possible mechanism for this effect could be through the modulation of attention by statistical learning (Batterink et al., 2015; Clement & Anderson, 2024; Cohen et al., 1990). By directing attention to certain features, statistical learning helps associate rewards with those features while suppressing others, effectively reducing dimensionality. In the case of the fruit example, animals might focus on the color and ignore size to quickly identify the most flavorful and nutritious options. However, despite this advantage, statistical relationships among cues, objects, and events may not always align with reward associations (e.g., for certain fruits, color may not indicate ripeness). This could cause statistical learning to bias reward learning towards features that are not particularly informative about reward outcomes. Nonetheless, regardless of whether statistical relationships and temporal regularities predict reward, attending to features with regularity while ignoring others can effectively reduce dimensionality.

Therefore, we hypothesized that detecting regularities in a certain feature of stimuli in the environment could impact feature-based learning from reward feedback on a separate set of stimuli that share this feature. To test this hypothesis, we designed a novel experimental paradigm involving two orthogonal tasks: a multidimensional reward-learning task to study more naturalistic reward learning, and a sequence-prediction task to engage statistical learning. During the sequence-prediction task, human participants were asked to predict the next stimulus in a sequence by identifying regularities within a feature. This was done to create temporal regularities in one feature (the feature with regularity) to focus attention on that feature, while keeping the sequence of the other feature random. During the multidimensional reward-learning task, participants learned the predictive value of a different set of stimuli by receiving reward feedback after choosing between pairs of stimuli. Importantly, each stimulus was associated with a reward probability that could be partially estimated using its features, with both features were equally informative in predicting the reward outcome. These two interleaved orthogonal tasks involved different stimulus sets that share the same feature dimensions (e.g., shape and pattern), enabling controlled manipulation of attention to a specific feature. We employed multiple analyses to examine how temporal regularities in the sequence-prediction task impact learning and choice behavior during reward learning. Finally, to uncover underlying computational and neural mechanisms, we developed various computational models and tested their ability to fit choice data on a trial-by-trial basis.

Our results demonstrate that temporal regularities in the sequence-prediction task influenced reward learning for a different set of stimuli that shared similar features. Specifically, the feature with temporal regularity was learned more strongly (showing larger learning rates), even though both features provided equal information about reward outcomes. However, we found little evidence that temporal regularities influenced decision making. Finally, the pattern of learning rates for chosen and unchosen options shows confirmation bias (Palminteri et al., 2017; Palminteri & Lebreton, 2022): learning rates for the chosen option were higher in rewarded trials than in unrewarded trials, and vice versa for the unchosen option. This bias was further amplified for the feature with temporal regularity. Overall, our findings support the hypothesis that statistical learning shapes reward learning by reducing dimensionality and further suggest that it may contribute to confirmation bias.

## Results

To investigate whether environmental regularities unrelated to reward can influence reward learning, we designed a novel experimental paradigm that involved two independent tasks: a multidimensional reward-learning task and a sequence-prediction task (**Fig. 1A**). During the multidimensional reward-learning task, human participants learned about the reward probabilities associated with 16 visual stimuli, each with two features (shape and pattern). The task consisted of two types of trials. In the choice trials, participants chose between pairs of stimuli and received probabilistic reward feedback from both the chosen and unchosen options, with the goal of maximizing their total reward. In value estimation trials, participants provided estimations of the reward probabilities associated with each stimulus.

**Figure 1.**
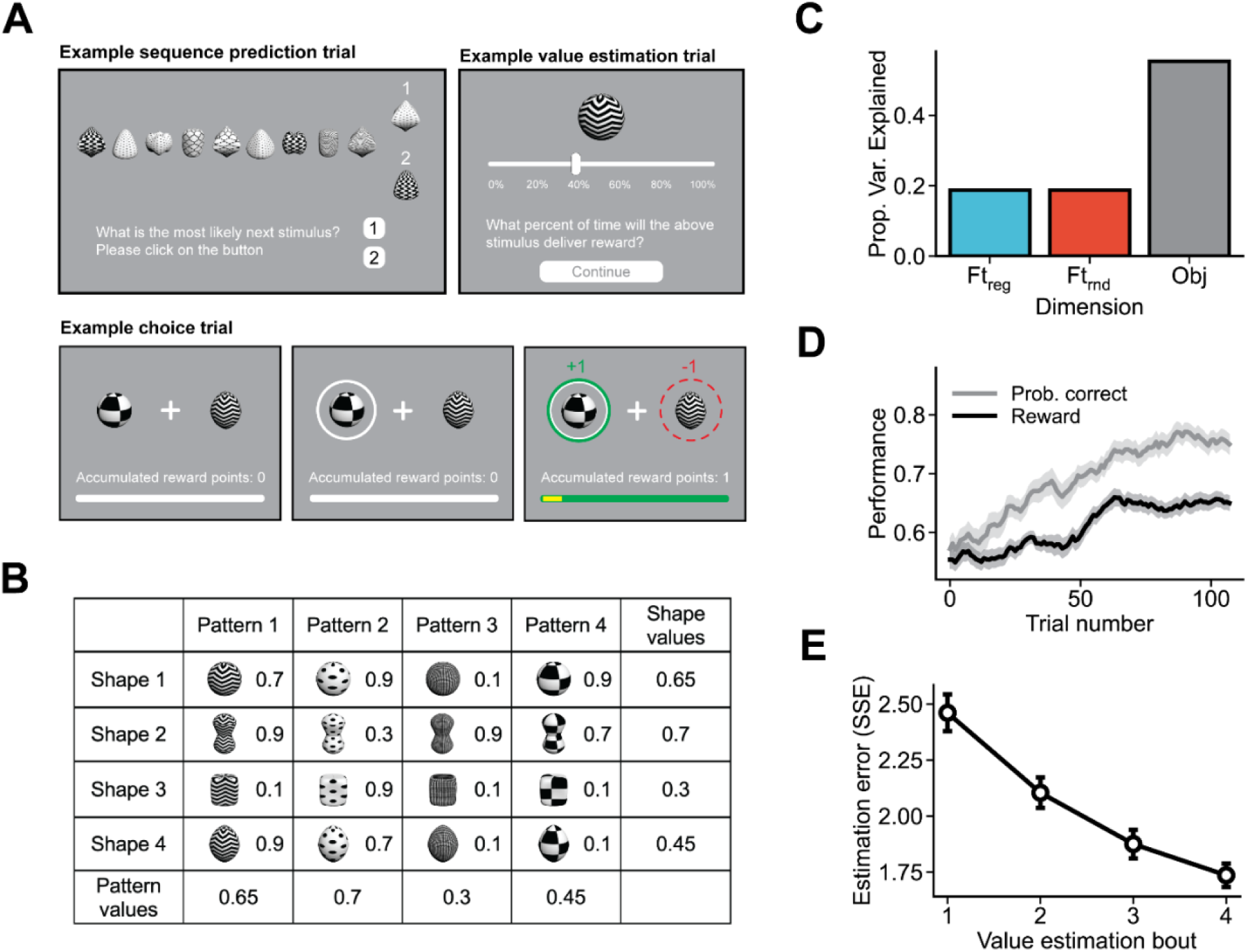
Experimental paradigm, the structure of the reward environment, and participants’ overall performance. (**A**) The experiment consisted of three types of trials. In sequence-prediction trials, participants were presented with a sequence of stimuli and asked to predict the next stimulus in the sequence. In choice trials, participants chose between a pair of stimuli and received reward feedback, which included the potential reward they would have received if they had selected the unchosen stimulus. In value-estimation trials, participants provided their estimates of the reward probabilities associated with all 16 stimuli. (**B**) Reward schedule. The table displays the reward probabilities associated with an example set of 16 stimuli used in the multidimensional learning task. Each stimulus was the combination of a shape and a pattern. Reward probabilities were symmetric for the two features. The shape and pattern values represent the mean reward probabilities for stimuli that share a common feature. (**C**) Variance decomposition of the reward schedule. Both features explained the same amount of variance in the reward probabilities, eliminating possible attentional bias due to differential informativeness of the two features. A substantial amount of variance was only explained by the combination of features, allowing us to disentangle feature-based and object-based learning. (**D**) Time course of learning during choice trials. The black and gray curves represent the average amount of reward obtained per trial and the proportion of trials in which the more rewarding stimulus was selected across all participants. The curves were smoothed using a moving average with a window of 20 trials. The shaded areas show the s.e.m. (**E**) Errors of participants’ value estimations as measured by the sum of squared error in each estimation bout. Error bars show the s.e.m.

The sequence-prediction task was specifically designed to create temporal regularities related to a certain feature of the stimuli used in the experiments. During each trial of the sequence-prediction task, participants were presented with a sequence of visual stimuli and were required to identify the next stimulus in the sequence. Critically, only one of the two features of the sequence of stimuli was repeated in doublets, triplets, or quadruplets. We refer to this predictable feature as the feature with regularity (denoted Ftreg). Detecting the feature with regularity and the sequence of its instance was necessary and sufficient for predicting upcoming stimuli and performing the sequence-prediction task correctly. In contrast, the other feature, which we refer to as the random feature (Ftrnd) was presented in random orders and thus could not be used to predict the upcoming stimulus. Crucially, while the stimuli used in the sequence-prediction task were also a combination of a shape and a pattern, they were distinct from the set of stimuli used in the reward-learning task, and information about temporal regularity was not useful for solving the reward-learning task.

Each participant initially performed 10-20 trials of the sequence-prediction task as practice. During the actual experiment, each participant went through 128 choice trials. One sequence-prediction trial was administered before every 8 choice trials, and a bout of estimation trials followed every 32 choice trials. This led to a total of 16 sequence prediction trials and 4 bouts of estimation trials during the main experiment (see **Methods** for more details). Sequence-prediction trials were interleaved with choice trials throughout the experiment to ensure the full impact of our manipulation was maintained.

Crucially, despite the difference in temporal regularity imposed during the sequence-prediction task, the reward schedule of the learning task was symmetric in terms of feature values, with both features being equally informative about the reward outcome (**Fig. 1B**). In addition, both features and their combination (the object identity) independently explained parts of the total variance in reward probabilities associated with the 16 stimuli used in the learning task (**Fig. 1C**). As a result, estimated reward probabilities based solely on feature value can be distinguished from actual reward probabilities associated with each object/stimulus (Farashahi et al., 2017). The Bayes-optimal feature-based learner should assign equal weights to both features (see Reward schedule and object-based vs. feature-based learning strategy in **Methods**), whereas a heuristic learner who is biased toward a specific feature, will assign a higher weight on that feature.

Despite the inherent difficulty of the reward-learning task, which involved 16 stimuli, along with the interleaved sequence-prediction task, the majority of participants successfully completed the experiment. Specifically, 57 out of the 78 total participants achieved a performance level above chance in both the learning and sequence-prediction tasks or their estimates of reward probability associated with each stimulus were significantly correlated with the assigned reward probabilities (see **Methods** for more details on exclusion criteria). The time course of participants’ performance in the choice trials indicated that learning continued throughout the experiment and only plateaued after 80 trials (**Fig. 1D**). In addition, the errors of their value estimations steadily decreased across the four bouts of value estimations trials (**Fig. 1E**).

### Effects of regularity manipulation on choice behavior and value estimations

To test whether temporal regularities in the environment modulated reward-based learning, we first analyzed participants’ choice behavior and value estimations. A previous study using a similar multidimensional learning task (without the interleaved sequence-detection task) suggested that the choice behavior of individuals who adopted feature-based learning can be captured by a weighted combination of feature values (Wang & Soltani, 2025). Therefore, we fit a mixed-effects logistic regression model using the log ratio of the true feature values and actual object values of the two options to predict participants’ choices (**Table S1**).

We found that participants’ choices were significantly influenced by the log of value ratio of the feature with regularity (*b*=1.20*, SE*=0.14, *z*=8.78, *p*=1.58×10^-18^), the log of value ratio of the random feature (*b*=0.78*, SE*=0.11*, z*=7.32, *p*=2.46×10^-13^), and to a much lesser extent, the log of value ratio associated with the stimulus itself (*b*=0.12, *SE*=0.04, *z*=3.09*, p*=0.002). This indicates that participants utilized a mixture of feature-based and object-based learning. These coefficients allowed us to illustrate the relationship between the probability of selecting one of two choice options and the log odds of the manipulated and random feature values for those options (Ftreg and Ftrnd, respectively; **Fig. 2A**). Importantly, the coefficients for the feature with regularity and random features were significantly different from each other (χ²(*1*)=7.34, *p*=0.007), suggesting that the values of the feature with regularity had a greater influence on choice behavior––even though, objectively, the two features were equally informative about reward. Correspondingly, the logistic curve for to the feature with regularity had a larger slope than that of the random feature (**Fig. 2A**).

**Figure 2.**
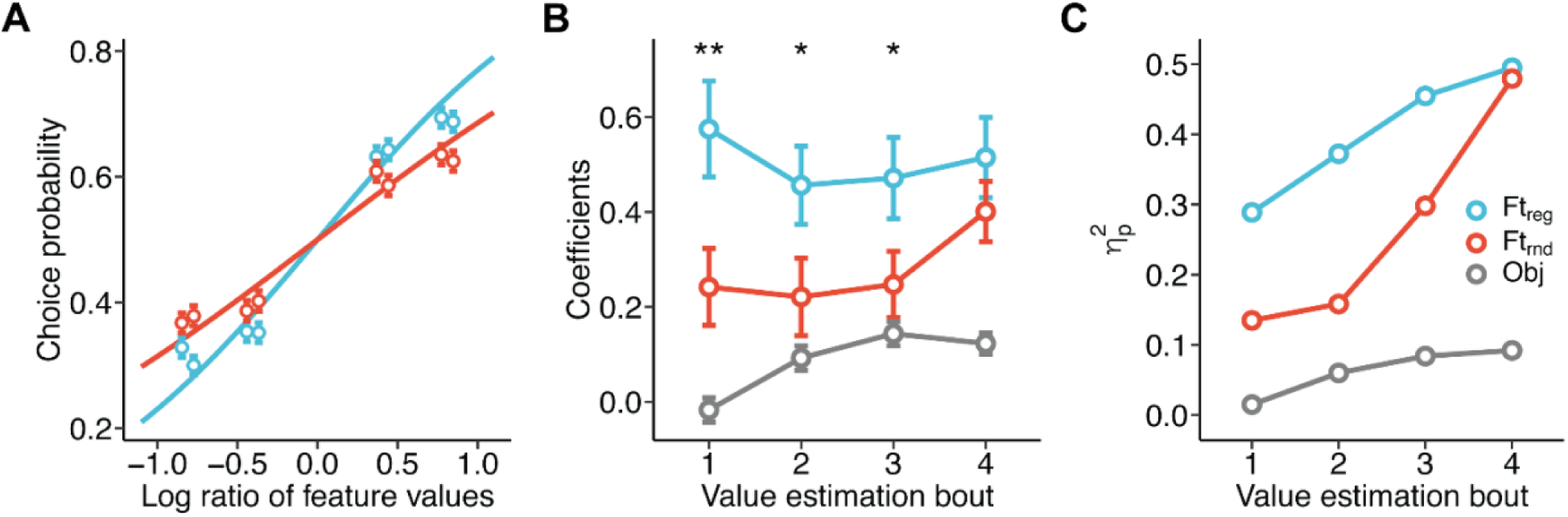
Choice and value estimation strategies during the reward-learning task. (**A**) Results of mixed-effects logistic regression using the log ratio of feature values to predict choice. Each point represents the choice probability associated with a specific log feature value ratio. Error bars show one s.e.m., and the lines show the best fit logistic curves. The slope for the feature with regularity (Ft_reg_) was significantly higher than that of the random feature (Ft_rnd_). (**B**) Results of a mixed-effects linear regression using feature and object values (Obj) to predict participants’ estimated stimuli values in four bouts of value estimation trials. Each point represents the fixed effects regression coefficient, with error bars indicating the standard error. Asterisks above the points indicate the significance of the contrast between the feature with regularity and random feature (\**p*<0.05; \*\**p*<0.01). Participants’ value estimations were initially better predicted by the values of the feature with regularity than by that of the random feature, and gradually became more consistent with the actual object/stimulus values. (**C**) Effect sizes of different fixed effects in the mixed-effects linear regression used to predict participants’ estimates based on feature and object identity. Each point represents the η_p_^2^ of each feature and the object. The feature with regularity accounted for more variance in the data compared to the random feature, suggesting that it had a more significant effect on participants’ estimation of values. This difference is reduced over time while the effect of object identity increases.

To gain a more fine-grained understanding of how participants computed and represented the reward probabilities of different stimuli, we examined their value estimates for all 16 stimuli, which were collected during four bouts of estimation trials that occurred throughout the experiment. For this analysis, we employed linear mixed-effects models that used the log odds of feature and object reward probabilities to predict participants’ estimations. We found that for first three estimation bouts, the coefficients of the feature with regularity were significantly higher than the coefficients of the random feature. The coefficient of the actual object values was not significantly greater than zero in the first bout but became significant in later bouts (**Fig. 2B**; **Table S2**).

A previous study involving a similar multidimensional learning task showed that feature values, defined as the average value of stimuli sharing a feature, may not fully reflect participants’ value estimations, as participants may not choose all stimuli sharing a feature with equal frequency (Wang & Soltani, 2025). Therefore, without assuming such feature values, we also predicted participants’ reward probability estimations using only the feature identities and their interactions as the independent variables (**Fig. 2C**; **Table S3**). By performing analysis of variance on the resulting linear mixed effects model, we found that for all trials, η_p_^2^ for the feature with regularity was higher than that of the random feature. Overall, this confirms that although both features had identical predictive values, participants placed greater weight on the values of the feature with regularity when estimating the values of stimuli. Moreover, in addition to rapidly acquiring the feature values, participants gradually learned the more accurate object values, as reflected in the increased influence of object identity on their estimations (**Fig. 2C**; **Table S3**). This is consistent with the previously reported transition from feature-based to object-based learning (Farashahi et al., 2017, 2020).

Overall, these results suggest that participants made choices and estimated values based on a weighted combination of feature values, with more weight given to the feature with regularity. To understand the emergence of this bias toward this feature, we next explored how choice on a given trial depended on choice and reward outcomes on the preceding trials.

### Effects of regularity manipulation on credit assignment and response to reward feedback

Previous studies on reward learning have demonstrated that both previous choice and reward outcomes can influence choice behavior (Barraclough et al., 2004; Kim et al., 2009; Witkowski et al., 2022). A general finding is that the reward outcome strongly influences future choices such that options that are rewarded are more likely to be selected again, whereas the absence of a reward typically leads to a switch from the previously chosen option. This pattern is commonly known as the ‘win-stay-lose-switch’ (WSLS) strategy and can be used to measure adjustments in learning and choice behavior (Trepka et al., 2021). Moreover, the selection of an option can increase the likelihood of choosing that same option in the future independently of reward outcome, a phenomenon commonly referred to as ‘choice autocorrelation’ (CA). Both WSLS and CA strategies yield predictions about subsequent choices based on choice and reward outcomes on a previous trial. These predicted choices can be used as regressors to fit the choice behavior of the participants in the upcoming trial (Katahira, 2018; Lau & Glimcher, 2005; Wang & Soltani, 2025). The resulting coefficients measure the extent to which the participants’ behavior can be described by each strategy.

Here, we expanded this approach in multiple ways to more thoroughly examine learning from multidimensional choice options. First, we defined WSLS and CA based on the individual features (i.e., Freg and Frnd) of the choice options in addition to their object identity to reveal both feature-based and object-based learning strategies. Second, to model counterfactual learning, we also included regressors for WSLS and CA based on the reward and choice histories associated with the *unchosen* options in the regression model. This was possible because reward feedback was provided for both chosen and unchosen stimuli in our study. Finally, to capture transitions from feature-based learning to mixed feature-based and object-based learning (Farashahi et al., 2017, 2020), we fit separate models to the trials in the first and second halves of the session. This led to twelve predictors (chosen/unchosen × WSLS/CA × feature with regularity/random feature/object) that were used in a mixed-effects logistic regression models to predict the choice behavior of participants during each half of the learning task (see Model-free analysis of choice and value estimations for more details). The resulting coefficients reflect how much participants utilized each strategy (**Fig. 3** and **Table S4**.)

**Figure 3.**
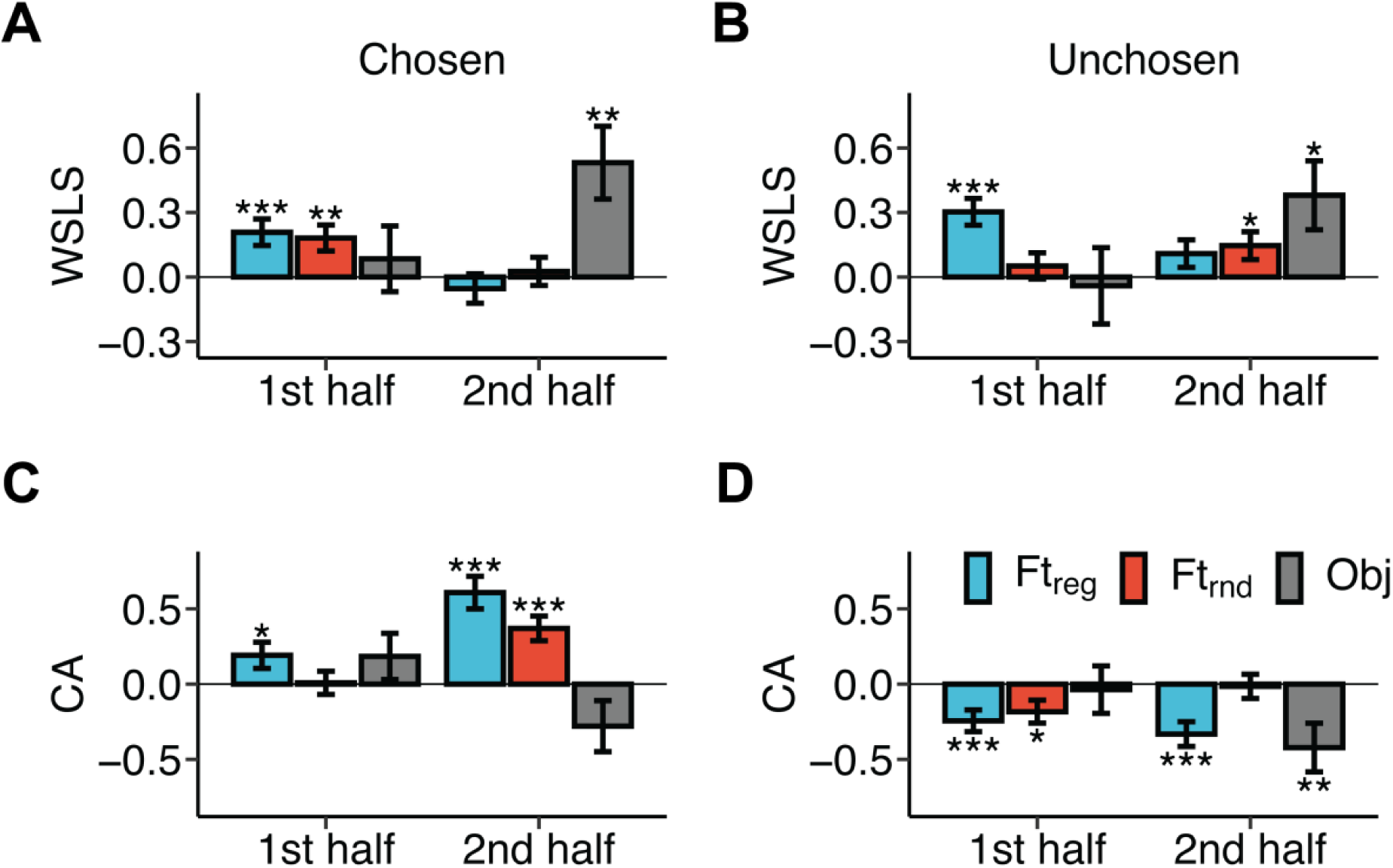
Contribution of different WSLS (win-stay-lose-switch) and CA (choice autocorrelation) strategies in participants’ choices, separately for the first and second halves of the experiment. (**A–B**) WSLS based on the chosen (A) and unchosen (B) options. (**C–D**) CA based on the chosen (C) and unchosen (D) options. Error bars represent 1 standard error, and asterisks indicate the significance level of the coefficient’s difference from zero (\**p* < 0.05, \*\**p* < 0.01, \*\*\**p* < 0.001). Ft_reg_, Ft_rnd_ and Obj denote the feature with regularity, random feature, and object, respectively.

Examining these coefficients provided several new insights into learning from multidimensional stimuli. Firstly, we observed a positive CA for the chosen option and a negative CA for the unchosen option (comparing **Fig. 3C** and **Fig. 3D**). A positive CA for the chosen option indicates that participants were inclined to select the previously chosen option or an option that shared one of its features. Conversely, a negative CA for the unchosen option implies that participants tended to avoid selecting either the previously unchosen option or an option sharing a feature with it, regardless of whether it was rewarded or not.

Secondly, we found a shift of learning strategies from feature-based to object-based learning (comparing the left and right sides of each panel in **Fig. 3**). In the first half of the trials, participants’ choices were only significantly influenced by reward and choice histories associated with the features of previous options, but not the combination of them (i.e., the object). In the second half of the study, the WSLS strategy based on the features of choice options was minimal, whereas CA based on option features was high. This indicates that by the second half of the experiment, participants have developed strong representations of feature values. At the same time, the reward and choice histories associated with the object (stimulus identity) significantly influenced their choice behavior. These findings suggest a shift from learning based on individual features to learning based on the stimulus/object (combination of features). Importantly, this shift was not due to object values being learned more slowly than feature values. Instead, in the first half of the trials, rewards were either not associated with the object or had minimal impact on subsequent choices.

Lastly, participants’ choices were more influenced by the reward and choice history of the feature with regularity (comparing blue and red bars of each panel in **Fig. 3**). In the first half of the experiment, the regression coefficients for the reward and choice histories of the feature with regularity were all significantly different from zero. In addition, the magnitudes of these coefficients were all larger than those for the random feature (**Fig. 3**). This effect was particularly evident in learning about the unchosen option, where participants showed significant WSLS behavior based on the feature with regularity (*b*=0.30, *SE*=0.06, *z*=4.87, *p=*1.13×10^-6^), but not based on the random feature (**Fig. 3B**). In the second half of the trials, participants’ choices were more consistently influenced by the feature with regularity, as indicated by a large positive CA for the chosen option and a large negative CA for the unchosen options (**Fig. 3C– D**).

Overall, these results provide compelling evidence that detecting temporal regularity during the sequence-learning task influence behavior during the reward-learning task. The observed changes in the behavior could occur via attentional modulations of reward learning and/or decision making. These effects occurred in tandem with other characteristics of human learning that have been observed in previous studies such as the transition from feature-based learning to object-based learning (Farashahi et al., 2017, 2020).

### Effects of regularity manipulation on feature-based learning

To uncover the computational and neural mechanisms underlying the observed patterns of learning and choice behavior, we next fit multiple reinforcement learning (RL) models to the choice behavior of participants. These models included an object-based learning model (Obj) and four feature-based learning models: a base feature-based model (Ft), and three feature-based models with attention modulating choice (FtAC), learning (FtAL), or both (FtACL). Our goals were to determine whether participants adopted feature-based or object-based learning and whether the manipulation of regularity modulated participants’ selective attention during reward learning, decision making, or both. We assumed that the effect of regularity manipulation remained constant throughout the session and modeled this effect as model parameters. As a result, the FtAC model has separate inverse temperatures for two features, the FtAL model has separate learning rates for the two features, and the FtACL has both (see Computational models in **Methods** for more details). Through model and parameter recovery analyses, we confirmed that these models could be distinguished from each other, and that their parameters could be reliably identified (**Fig. S1**).

Using the Watanabe-Akaike information criterion (WAIC) to measure goodness-of-fit, we found that participants’ behavior was best described by the feature-based learning model FtAL, in which attention modulated learning only (**Fig. 4**). Interestingly, the model in which attention modulated both choice and learning (FtACL) did not improve the fit of choice data compared to the best model (*ΔWAIC* = 0.93 ± 2.29). Moreover, the model in which attention modulated only decision making (FtAC) showed a significantly lower goodness-of-fit compared to the best model (*ΔWAIC* = 11.77 ± 5.68), and similarly, the base model with no attentional modulation (Ft) also exhibited reduced fit quality (*ΔWAIC* = 34.89 ± 11.79). The object-based model (Obj) had the worst fit overall (*ΔWAIC* = 109.60 ± 35.10).

**Figure 4.**
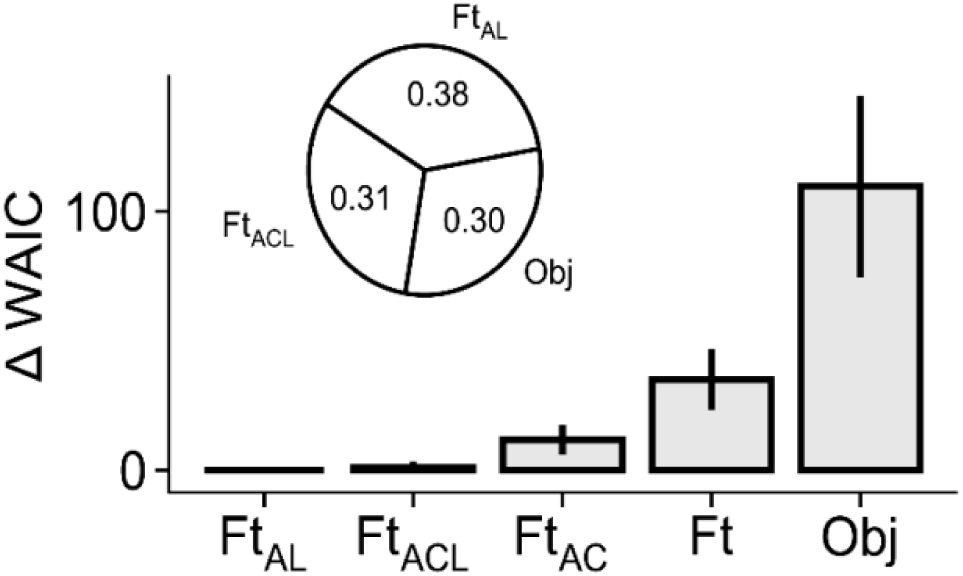
Participants’ choice behavior was best explained by a feature-based model in which attention led to differential learning of the two features. The plot compares the goodness-of-fit of different models, measured by the difference in WAIC between each model and the best-fitting model, Ft_AL_ (feature-based learning with attention modulating learning). The pie chart shows the stacking weights calculated using WAIC. Model abbreviations: Ft_ACL_ (feature-based learning with attention modulating choice and learning), Ft_AC_ (feature-based learning with attention modulating choice), Ft (base feature-based learning), Obj (object-based learning). Error bars show 1 s.e.m. Feature-based learning models that included attentional modulation of learning were assigned larger weights than models without such modulation. Object-based learning also has substantial weight despite having overall worse fit, suggesting its importance in accounting for participants’ choice behavior.

We also examined the stacking weights, which indicate how different models can be combined to maximize predictive performance (Yao et al., 2018). This approach does not assume that the true model is among the included models and can identify models that, despite having a worse overall WAIC, are still useful for predicting behavior. Indeed, we found substantial weights not only for the best fitting model (FtAL) and feature-based learning with attentional modulation of both learning and choice (FtACL), but also for the object-based learning model, with stacking weights of 0.38, 0.31, and 0.30 respectively (**Fig. 4** inset). Overall, these results demonstrate that participants employed a combination of feature-based and object-based learning strategies, with attention primarily modulating the learning of feature values rather than their integration during decision making.

To further pinpoint the effect of regularity on value learning, we next compared the estimated learning rates for the two features in the best-fitting model (**Fig. 5A–D**). We found that the learning rates for the feature with regularity were generally higher than those for the random feature (*median difference*=0.022, *probability of direction*=0.99, *95% HPDI:* [0.0040, 0.043]). By comparing each pair of learning rates for the two features, the most pronounced difference was observed in the learning rate on the unchosen option when it was not rewarded (*median difference=*0.061*, probability of direction=*0.97, 95% *HPDI:* [-0.0013, 0.13]; **Fig. 5E–H**) (see Parameter estimation and hypothesis testing in **Methods** for more details). This finding further corroborates the results from mixed-effect logistic regression analysis on choice in the previous section, which showed the effect of regularity manipulation on learning was the strongest when learning about the unchosen option (**Fig. 3B**). Notably, simulations across a wide range of learning rate biases revealed that assigning a larger learning rate to either feature through selective attention resulted in poorer performance (**Fig. S2**). This is expected, as the sequence-prediction task was orthogonal to the reward-learning task. Nonetheless, it underscores the effectiveness of the regularity manipulation.

**Figure 5.**
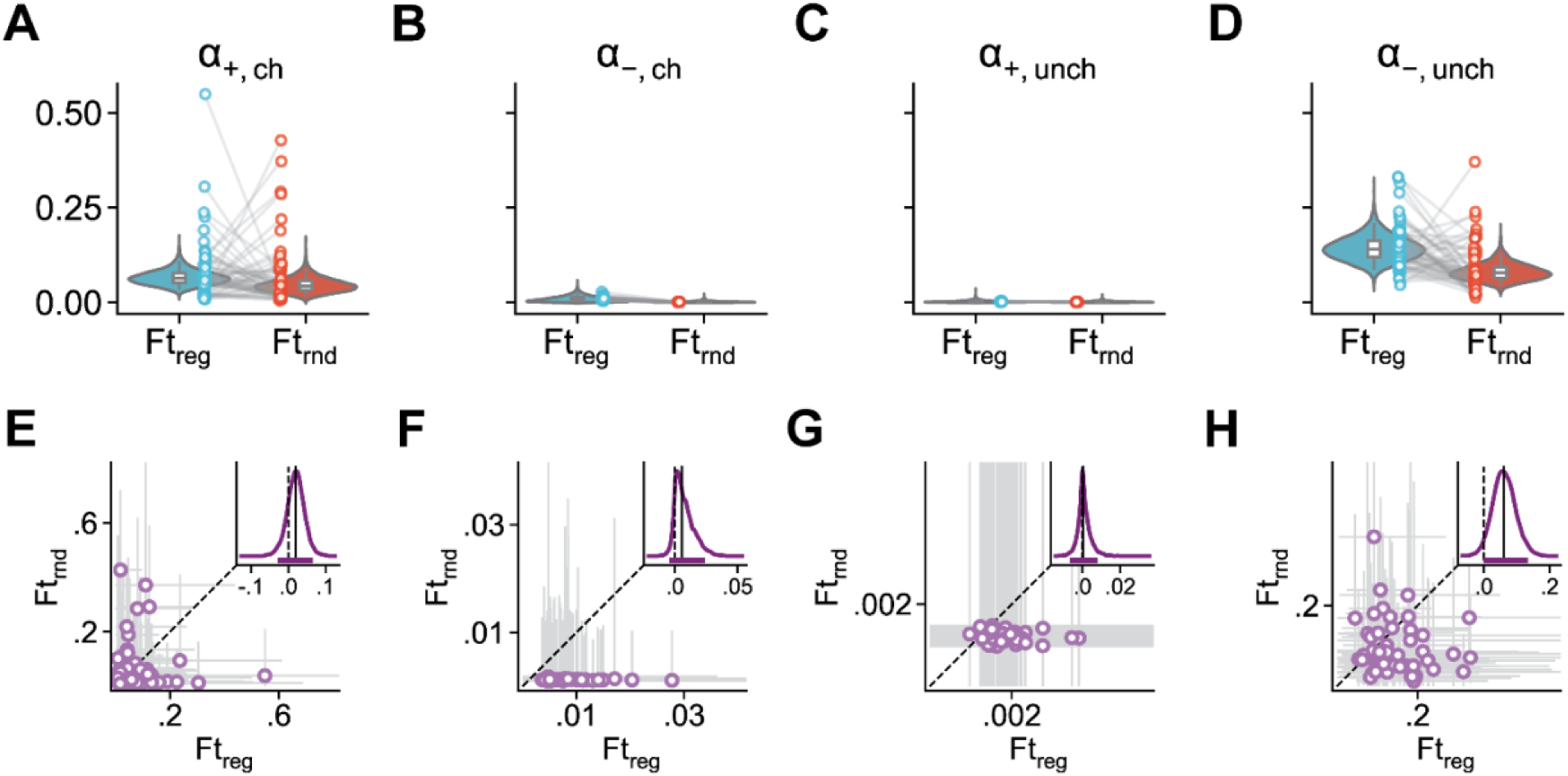
Sequence-prediction task led to modulations of learning during the reward-learning task as reflected in higher learning rates and more pronounced confirmation bias for the feature with regularity. (**A–D**) Evidence of strong confirmation bias in feature-based learning. Violin plots show posterior distributions of group-level learning rates for chosen (ch) and unchosen (unch) options and separately for rewarded (*α*_+_) and unrewarded (*α*_−_) trials. Each point shows the learning rates for individual participant and whiskers of the box plot show 95% the HPDI. (**E–H**) Evidence of regularity detection influencing the learning rates, particularly about unchosen and unrewarded options. Scatter plots show the posterior distributions of individual participants’ learning rates. Dots show the median and grey bars show the 95% HPDI. The insets plot the posterior distribution of the difference between the group-level learning rates for the feature with regularity and the random feature. Vertical solid line indicates the median. Dotted line indicates zero. The purple band shows the 95% HPDI.

### Enhanced confirmation bias due to regularity manipulation

In addition to an overall increase for the feature with regularity, the estimated learning rates showed a pattern consistent with confirmation bias in learning. Specifically, the learning rates for the chosen option were higher in rewarded trials than in unrewarded trials (feature with regularity: *median difference=*0.055*, probability of direction*>0.999, 95% *HPDI*: [0.018, 0.099]; random feature: *median difference=*0.042, *probability of direction*>0.999, 95% *HPDI*: [0.011, 0.082]; **Fig. 5A, B**), a phenomenon referred to as the positivity bias in reward learning (Palminteri et al., 2017; Palminteri & Lebreton, 2022). In contrast, the learning rates for the unchosen option were larger in unrewarded than in rewarded trials (feature with regularity: *median difference*=-0.14, *probability of direction*>0.999, 95% HPDI: [-0.20, -0.075]; random feature: *median difference=*-0.076*, probability of direction*>0.999, 95% *HDPI*: [-0.12, -0.040]; **Fig. 5C, D**), a pattern termed the negativity bias in reward learning. Notably, simulations across a wide range of confirmation levels revealed a significant benefit of confirmation bias for performance in the current task (**Fig. S2**).

Interestingly, the overall confirmation bias––the average of positivity biases when learning about chosen options and negativity biases when learning about unchosen options––was more pronounced for the manipulated than for the random feature (*median difference*=0.036, *probability of direction*=0.98, *95% HPDI:* [0.00091, 0.074]). Individually, this difference was significant for the negativity bias (*median difference* =0.060*, probability of direction*=0.97, 95% *HPDI*: [-0.0025, 0.13]), and not for the positivity bias (*median difference*=0.013, *probability of direction*=0.71, *95% HPDI*: [-0.039, 0.059]). Consistently, the negativity bias was larger than the positivity bias, though this effect was only significant for the feature with regularity (manipulated: *median difference*=0.083, *probability of direction*=0.98, 95% *HPDI*: [0.013, 0.16]; non-manipulated: *median difference*=0.034, *probability of direction*=0.88, 95% *HPDI*: [-0.024, 0.094]). Overall, these results indicate that attentional modulation, triggered by regularity detection, can enhance confirmation bias in reward learning.

To confirm that regularity manipulation has distinct effects on the learning rates for chosen and unchosen options depending on the reward outcomes, we performed additional control analyses using simplified versions of the full FtAL model. In these models, the effect of attentional modulation was either considered uniform across all learning rates (the Ftunif model), or coupled specifically as the confirmatory learning rate (*α*_+,*ch*_ = *α*_−,*unch*_) and the disconfirmatory learning rate (*α*_−,*ch*_ = *α*_+,*unch*_) in the Ftcoupled model. We found that both simplifications resulted in a worse fit compared to the full FtAL model (*ΔWAIC* (*Ft*_*coupled*_ − *Ft*_*unif*_) = 13.65 ± 7.50; *ΔWAIC* (*Ft*_*coupled*_ − *Ft*_*unif*_ ) = 38.01 ± 11.58; **Fig. S3A**). This demonstrates that regularity had distinct effects on confirmatory and disconfirmatory learning rates depending on the reward outcomes.

Overall, our findings align with the confirmation bias observed in reward learning (Palminteri & Lebreton, 2022) and importantly, extend this bias to feature-based learning, where we discovered a larger negativity bias compared to positivity bias. Additionally, they demonstrate that attention evoked by regularity detection can enhance confirmation bias, particularly by increasing the learning rate for counterfactual and unrewarded outcomes.

### Changes in learning strategy and effects of regularity manipulation over time

To identify possible changes in learning strategy and the effects of regularity manipulation over time, we first analyzed choice data from the first half of the experiment using the same models as in the previous section. Consistent with the results based on all data, the FtAL model provided the best fit for participants’ choice behavior in the first half of the experiment (**Fig. S3B**). However, analysis of the stacking weights further revealed that object-based learning was not useful for predicting behavior in the first half of the trials (compare **Fig. S3B** and **Fig. 5**), indicating that participants relied more heavily on feature-based learning at the beginning of the experiment.

Nonetheless, confirmation bias was present in the first half of the experiment (**Fig. S4A–D**). Moreover, the influence of regularity on the learning rates was more pronounced in the first half of the experiment than across the entire experiment, particularly for the learning rate of unrewarded, unchosen options (*median difference=*0.13*, probability of direction=*0.999, 95% *HPDI:* [0.057, 0.23]) (**Fig. S4E–H**).

To better assess how feature-based and object-based learning strategies competed against each other over time, we examined the trial-wise differences in WAIC between the best-fitting feature-based model (FtAL) and the object-based model (**Fig. 6A**). By fitting a linear mixed-effects model with trial number to predict the trial-wise difference in WAIC (between the feature-based and object-based model) for each participant, we found that the gap significantly decreased with time (*b*=-0.04*, SE*=0.01*, t*(260.5)=-2.82*, p*=0.005). This indicates that participants’ behavior became more consistent with the object-based model later on in the experiment. The intercept was significantly larger than zero, confirming the fact that the feature-based model had significantly better fit at the beginning of the experiment (*b=*0.04*, SE=*0.007*, t*(7239)=5.11, *p*=3.34×10^-7^).

**Figure 6.**
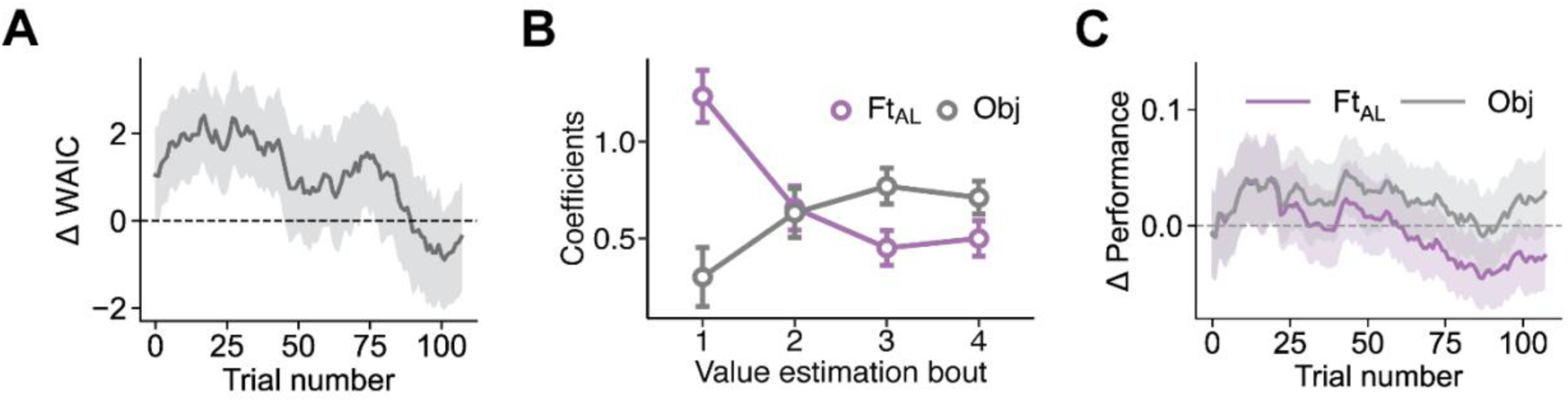
Gradual transition from feature-based to object-based learning. (**A**) Difference between the goodness-of-fit of the best feature-based (Ft_AL_ ) and the object-based model (Obj) over time, measured by ΔWAIC. The shaded area represents one standard error. Initially, the feature-based learning model fits behavior better, but the goodness-of-fit of the two models becomes similar by the end of the experiment. (**B**) Influence of feature-based and object-based learning on value estimates. Plot shows the results of linear mixed-effects regression predicting value estimates in each bout of estimation trials using reward values learned by the Ft_AL_ and Obj models. Each point represents regression coefficients, and error bars represent one standard error. Early on, value estimates were more strongly influenced by feature values, but later in the experiment, they became better predicted by object values. (**C**) Comparison of the participants’ learning curves and that of the Ft_AL_ and Obj model. The shaded area represents the 95% HPDI. The Ft_AL_ model performed similarly to participants initially but underperformed later in the session. In contrast, the Obj model outperformed participants overall, yet more closely matched their performance in the later trials.

In addition, we also used linear mixed-effects models to predict participants’ value estimates during each bout of estimation trials based on the values estimated by the feature-based and object-based models (**Fig. 6B; Table S5**). This analysis revealed that participants’ value estimates were strongly feature-based in the first bout of trials, as the values learned by the best-fitting feature-based model were assigned a larger coefficient. However, later in the experiment, feature and object values had similar influence, with the coefficients on object value being slightly higher. This suggests that participants dynamically arbitrated between feature-based and object-based learning models to control their behavior (Farashahi et al., 2017, 2020; S. W. Lee et al., 2014).

Finally, we validated the models by simulating the best-fitting feature-based and object-based models using their estimated parameters. We then examined whether the performance of these models, in terms of the probability of selecting the better option on each trial, matched that of human participants. By calculating the difference between the empirical learning curve and the learning curves of each model, we found both model’s simulated performance closely matched the empirical performance (**Fig. 6C**). At the beginning of the experiment, however, the performance of the feature-based model aligned with that of the participants, whereas the performance of the object-based model remained above that of the participants. On the other hand, toward the end of the experiment, the performance of the feature-based model dropped below that of the participants, whereas the performance of the object-based model stayed closer to theirs. We found consistent results when we conducted similar analyses separately for participants whose choice behavior was better explained by feature-based or object-based learning models. Importantly, each group of participants’ performances were better matched by the model that best described their respective choice behavior (**Fig. S5**).

This result was further confirmed by model-free analyses of the simulated choices of the FtAL and object-based models. The FtAL made choices by taking a weighted combination of feature values, with a higher weight placed on the feature with regularity (*χ^2^*(1)=15.21, *p*=9.60×10^-5^; **Fig. S6A**), consistent with the empirical data (**Fig. 2A**). Similarly, the values assigned to each stimulus/object by the FtAL model were consistent with a weighted sum of feature values, with a higher weight placed on the feature with regularity across all four estimation bouts (**Fig. S6B**).

This was more consistent with empirical value estimations in earlier trials (**Fig. 2B**). On the other hand, the values learned by the object-based model were more consistent with the stimulus/object values without a significant bias towards either feature value (**Fig. S6C**), similar to characteristics of empirical value estimations in later trials (**Fig. 2B**). The empirical credit assignment behavior (**Fig. 3**) was also similar to a mixture of the FtAL model (**Fig. S7A–D**) and object-based model (**Fig. S7E–H**). Finally, the FtAL accounted for feature-based learning biases toward the feature with regularity. This effect was evident in the early trials and persisted into the later trials. In contrast, the object-based model captured object-based learning, which emerged later in the experiment.

Together, these results demonstrate that, consistent with previous studies (Farashahi et al., 2017, 2020; Wang & Soltani, 2025), participants dynamically transitioned from learning about features to learning about objects. Moreover, modulations of learning and confirmation bias became evident early in the experiment when participants’ behavior was better described by feature-based models. This suggests that learning regularities influences multidimensional reward learning, highlighting the impact of statistical learning on reward learning beyond just mitigating the curse of dimensionality.

## Discussion

Using a novel experimental paradigm involving two orthogonal tasks combined with multiple computational approaches, we investigated the impact of environmental regularities on reward learning. We found that human participants leveraged regularities about features from a set of stimuli presented during a sequence-prediction task to adjust their reward learning about another set of stimuli that contain similar features. Both choices and estimations made by participants during the learning task were more heavily influenced by the objective values associated with the feature with regularity. This occurred despite the regularity not being informative about reward outcomes. Furthermore, through multiple model-free and model-based analyses, we established that these effects are more consistent with attentional modulations of learning rather than decision making. The learning rates for the feature with regularity were higher, particularly when learning from the chosen, rewarded options and from unchosen, unrewarded options.

This suggests that environmental regularities can intensify the confirmation bias observed in reward learning, perhaps due to feature-based attention driven by detection of regularity.

Consistently, previous studies have demonstrated that attentional mechanisms are both used and necessary for statistical learning (Batterink et al., 2015; Cohen et al., 1990), and that environmental regularities can modulate attention spontaneously, regardless of goal relevance or intrinsic salience (Kim & Anderson, 2021; Le Pelley et al., 2022; Ogden et al., 2023; Stankevich & Geng, 2014; Zhao et al., 2013). These attentional modulations can influence various cognitive processes including learning and decision making. Therefore, the observed difference in learning rates in our experiment may be mediated by attentional effects on reward learning, though differently from effects reported previously. Specifically, previous studies have shown that learning and decision making are modulated by attentional weights, which are determined by the value of features (Leong et al., 2017; Wang & Soltani, 2025). In our study, we controlled for the informativeness of two features to ensure that any observed differences in feature-based learning were due to regularities rather than reward value. Furthermore, to prevent contamination of reward value from the learning task influencing feature-based selection in the sequence-prediction task, we employed two completely different sets of stimuli for each task. Therefore, our results suggest that knowledge acquired through both reward feedback and statistical learning can shape future reward learning.

We found that attention evoked by environmental regularities primarily affects learning, rather than decision making. In contrast, prior studies on value-based attention have shown its impact on both learning and decision making (Leong et al., 2017; Niv et al., 2015), whereas others suggested it influences only the learning process, not decision making (Wang & Soltani, 2025). This difference may stem from variations in the experimental paradigms used or from the inherent challenges in detecting attentional modulation within RL models. Most previous studies used models in which only the value of the chosen option was updated based on reward feedback while allowing the value of the unchosen option to decay. In our study, however, feedback was presented for both the chosen and unchosen options allowing us to capture the differential effect of attention on updating the values of both chosen and unchosen options. This could favor the specific effect of attention during learning we found. However, our model recovery results suggest a bias towards detecting attentional effects on learning rather than on decision making.

This indicates that there could be undetected effects of attention on choice.

Our finding that regularity manipulation strongly influences reward learning through attentional mechanisms suggests that attention driven by statistical learning could be as powerful, if not more so, in shaping behavior than attention driven by predictive reward values. This is consistent with the literature on the effects of reward on perception and working memory (Hickey et al., 2010b, 2010a, 2014; Infanti et al., 2015), and with statistical learning literature (Jiang et al., 2015; Rogers et al., 2016). For example, a previous study that compared the effects of goal-driven attention, statistical learning, and monetary reward found that attention driven by statistical learning significantly affected search times, whereas the impact of reward-driven attention was negligible (Jiang et al., 2015).

Studies of confirmation bias in reinforcement learning have shown a stronger learning from confirmatory rather than disconfirmatory evidence in both human and non-human animals (Doll et al., 2011; Lefebvre et al., 2022; Palminteri et al., 2017). This bias has significant implications for reinforcement learning, potentially leading to both biased and optimal learning outcomes depending on the reward environment (Palminteri & Lebreton, 2022). Here, we find evidence for a novel extension of confirmation bias to feature-based learning, suggesting that the underlying mechanisms (attentional modulation) may be generalizable. This bias was present early in the experiment and was not affected by the transition from feature-based to object-based learning.

Interestingly, we observed a stronger negativity bias than positivity bias. A strong negativity bias can effectively reduce the value of less rewarding (and less frequently chosen) options to zero, thereby removing them from the choice set. Nonetheless, the contrast between positivity and negativity bias suggests that mechanisms underlying learning from factual versus counterfactual outcomes may be fundamentally different (Boorman et al., 2011; D. Lee et al., 2005).

Therefore, our results demonstrate that learning regularities about features can significantly influence feature-based reward learning and help mitigate the curse of dimensionality in two ways: reducing the number of relevant features through selective attention, and by eliminating less rewarding options from the choice set through negativity bias, which suppresses their overall value. These novel interactions between statistical learning and reward-based learning have interesting implications for learning behaviors, particularly in naturalistic settings.

Given the observed effects of regularity on reward learning, one might question whether the incorporation of regularities ultimately benefits learning or could lead to maladaptive behavior. In our study, we ensured that statistical relationships in the environment were not predictive of reward outcomes and therefore, meaning that stronger learning from the feature with regularity was not beneficial. Similarly, if the feature with regularity is not informative about reward, incorporating regularity into learning could be maladaptive. In many cases, however, learning the statistical structure of reward outcomes can improve reward learning by providing predictions in addition to enhancing simple reward associations (Atilgan et al., 2022; Fritsche et al., 2023; Gibson, 1969; Klein-Flügge et al., 2019; Soltani & Koechlin, 2022; Woo et al., 2023).

Moreover, acquiring statistical knowledge of the environment can facilitate faster learning of new or modified sequences compared to the rigid stimulus-stimulus associations typically seen in purely RL models (Klein-Flügge et al., 2019). Therefore, while detecting features with regularities and using them for learning can mitigate the curse of dimensionality, these regularities may also provide information about environmental changes related to finding food (e.g., day-night cycles), shelter (e.g., seasonal changes), mates (e.g., hormonal cycles), and more. Consequently, detecting these regularities and utilizing them for learning can ultimately help survival by facilitating learning in high-dimensional reward environments and reducing their dimensionality. Together, the observed interplay between statistical learning, attention, and reward learning has important implications for understanding learning in naturalistic settings, in both biological and artificial systems.

## Methods

### Participants

78 participants were recruited from Dartmouth College (53 females, age 18-32, mean age = 19.65). The sample size in this study was determined based on previous studies that used a similar approach, ensuring consistency and comparability of results (Farashahi et al., 2017; Farashahi & Soltani, 2021Wang & Soltani, 2023). We excluded 18 participants who did not perform significantly higher than chance level (50%) in the second half of the choice trials according to a binomial test (α=0.05) or whose value estimations did not correlate significantly with the actual object/stimulus values (using Spearman correlation, α=0.05) in the final two estimation bouts. We also excluded two participants who did not perform significantly higher than chance during the sequence-prediction task. Finally, one participant was excluded who failed to complete the experiment due to a hardware malfunction. This led to the inclusion of 57 participants in the analysis. Participants were compensated based on their performance (up to $10) in addition to a baseline compensation (either $10 or 1 T-point which is Dartmouth College credit compensations for some courses within the Psychological and Brain Sciences Department). During the experiment, accumulated reward points were displayed on the screen using a reward bar after each reward feedback, with every 10 reward points earning them 1 dollar. The experiment was approved by Dartmouth college Institutional Review Board, and a written consent form was obtained from each participant at the beginning of the experiment. This study was not preregistered. We report how we determined our sample size, all data exclusions, all manipulations, and all measures in the study.

### Visual stimuli

The experiment was created using jsPsych (Leeuw et al., 2023) and presented within a web browser of the computer in a controlled experiment room. We created the stimuli from a set of “Quaddles” all in grayscale (Watson et al., 2019). Each stimulus was composed of two features (shape and pattern) from eight unique shapes and eight patterns. For each participant, two distinct sets of 16 stimuli were created from four shapes and four patterns each, ensuring no object in one set had the same shape or pattern as any object in the other set. One set was used for the multidimensional learning task, and the other for the sequence-prediction task, with assignments counterbalanced across participants. This separation aimed to eliminate any bias from the effect of temporal regularities in the sequence-prediction task on participants’ expectations about upcoming features during the reward-learning task.

### Reward schedule and object-based vs. feature-based learning strategy

We designed a specific reward schedule to allow adoption of different learning strategies while ensuring that two features of stimuli/object are equally informative about reward outcomes (**Fig. 1B-C**). More specifically, an object-based learner chooses between the two choice options (stimuli) by estimating the reward probabilities of the two options directly. In contrast, a feature-based learner estimates the reward probabilities of the features separately and combines them to estimate reward probability for each stimulus/object. The feature values are defined as the average reward probabilities of stimuli/objects that contain a given feature (i.e., the shape and pattern values in **Fig. 1B**):

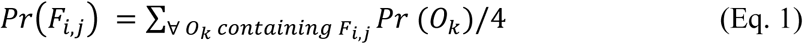

where *Fi,j* denotes the j-th instance of the i-th feature, *Ok* denotes the k-th object, and *Pr* denotes the probability of reward. We refer to these as the *true* feature values to differentiate them from the subjective values learned by reinforcement learning models (see **Computational models** below).

Using the Naive Bayes equation (Farashahi et al., 2017; Hunt et al., 2014), one can combine the feature values to estimate the stimulus/object values:

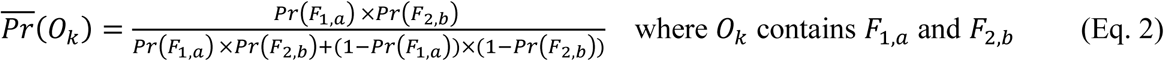

Taking the logit transformation of both sides reveals that the log odds of reward for each object can be estimated by linearly combining the log odds of reward for each feature:

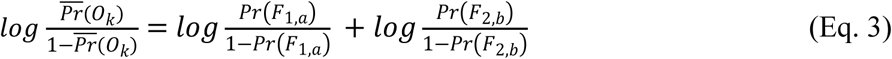

However, an agent may calculate a weighted combination of feature values to estimate the reward probabilities associated with each stimulus or object:

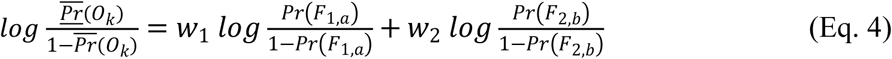

where *w1* and *w2* are the weights of the two features. Estimation of object values based on weighted average of feature values will differ from the actual reward probabilities of objects, leading to estimation errors that allow us to disentangle feature-based and object-based strategies. In our experiment, each instance of feature 1 had a corresponding instance in feature 2 with the same reward probability, meaning both features were equally informative about the reward outcome (**Fig. 1B**). This symmetric reward schedule was implemented to prevent value-based attentional biases toward any particular feature (Farashahi et al., 2017; Farashahi & Soltani, 2021; Leong et al., 2017; Niv et al., 2015; Wang & Soltani, 2023).

### Experimental procedure

The experiment consisted of two tasks and three types of trials: the sequence-prediction task that involved only one type of trials, and the multidimensional learning task that involved choice and estimation trials (**Fig. 1A**). The sequence-prediction task was designed to introduce temporal regularity related to a specific feature of the stimuli, each of which contained two features: shape and pattern. In each trial of this task, participants were presented with a sequence of visual stimuli and were required to identify the next stimulus in the sequence. The sequence of presented stimuli contained a repeating pattern (doublets, triplets, or quadruplets) for one of the two features, making it predictive of the next stimulus. The other feature appeared in a random order. We refer to the predictable and random features as the feature with regularity (Ftreg) and the random feature (Ftrnd), respectively. Temporal regularity of this kind has been shown to guide feature-based learning towards the more predictable feature (Chun & Jiang, 1999; Zhao et al., 2013).

The length of the presented stimuli (sequence) varied between seven to eleven stimuli/objects. Following the presentation of a sequence, two choice options were presented (**Fig. 1A**). After making a choice, participants received feedback indicating whether their choice was correct, along with the correct sequence (i.e., no reward feedback was provided). Performance in this task did not affect the monetary reward that participants would receive. Each participant completed up to 20 practice sequence-prediction trials at the start of the experiment. After the first five practice trials, those who achieved five consecutive correct predictions could immediately proceed to the main experiment. During the main experiment, one sequence-prediction trial occurred before every set of eight choice trials.

During choice trials of the learning task, participants had to choose between a pair of stimuli/objects using the F and J keys on a keyboard. After choosing between the two options, a white circle would appear around the chosen stimulus/object indicating the choice of the participant. Feedback for both stimuli were then presented (green circle and +1 for the rewarded and red circle and -1 for unrewarded option). An accumulated reward point representing the points the participant had earned would be shown at the bottom of the screen and after every 10 reward points followed by a message “You earned one dollar!” and then would be reset again. For the choice trials, each stimulus/object was assigned a reward probability unknown to the participants and had to be learned through trial and error. The relationship between the perceptual properties of the stimuli and the reward probability was randomly shuffled across participants. We selected a subset of 32 pairs of stimuli/objects to present as alternative choice options during choice trials (**Fig. 1B**). Each object with pattern 1 or 2 and shape 1 or 2 was paired with an object with pattern 3 or 4 and shape 3 or 4, and each object with pattern 3 or 4 and shape 1 or 2 was paired with an object with pattern 1 or 2 and shape 3 or 4. These pairs were selected to lower the difficulty of the task, while still allowing us to detect different learning strategies. The experiment consisted of a total of 128 choice trials. Each pair of choice options was presented in varying order arrangements during the first 64 trials and again in the last 64 trials, with the orders randomized across participants.

In value estimation trials of the learning task, we asked participants to indicate their estimation of the reward probability of each stimulus/object using a sliding scale (**Fig. 1A**). In each bout of estimation trials, participants were shown each of the 16 stimuli/objects in random order, one in each trial, and were asked to estimate the reward probability of the presented object with a slider ranging between 0-100%. There was no feedback in the estimation trials. One bout of value estimation trials happened after every 32 choice trials.

Critically, we made the sequence-prediction and learning tasks orthogonal in two ways. First, we separated the reward and regularities by not providing reward during the sequence-prediction task. Second, we used different sets of stimuli/objects in the two tasks so that the learning about stimuli/objects in one task did not influence the learning about stimuli/objects in the other task.

Finally, to encourage prompt responses and engagement, we set a time limit of fifteen seconds for the sequence-prediction and value-estimation trials, and eight seconds for choice trials.

Warnings were displayed at ten seconds, seven seconds, and five seconds before the time limit in sequence-prediction trials, value-estimation trials, and choice trials, respectively. After reaching the time limit, the trial was skipped with a warning. The experiment would be terminated if a participant skipped four trials.

### Model-free analysis of choice and value estimations

To explore how participants adjusted their choice behavior based on reward and choice history, we fit generalized linear mixed effects models to predict participants trial-by-trial choices using the choice and reward information from the previous trial:

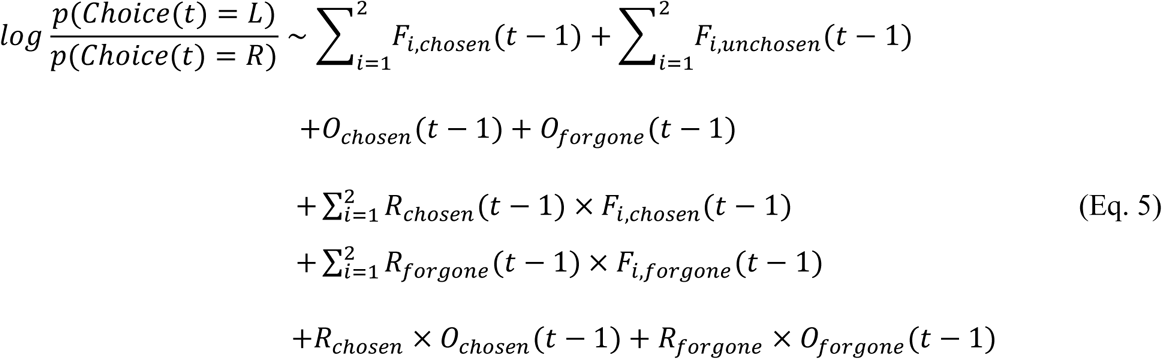

𝐹_𝑖,*ch*𝑜𝑠𝑒𝑛_ is equal to +1 if the left option shared the i-th feature with the previously chosen option, -1 if the right option shared the i-th feature with the previously chosen option, and 0 otherwise. 𝐹_𝑖,*unch*𝑜𝑠𝑒𝑛_ is defined analogously but based on the previously unchosen option. 𝑂_*ch*𝑜𝑠𝑒𝑛_ and 𝑂_*unch*𝑜𝑠𝑒𝑛_ are defined analogously depending on whether the choice option (stimulus/object) on a given trial is the same as the stimuli/object on the previous trial. 𝑅_*ch*𝑜𝑠𝑒𝑛_ and 𝑅_𝑓𝑜𝑟𝑔𝑜𝑛𝑒_ are the rewards associated with the chosen and unchosen options (rewarded: +1; unrewarded: -1).

To examine participants’ value representation more directly, we fit linear mixed effects models to predict participants’ value estimations from the feature and object values. Both value estimations and the feature and object values were put through a logit transformation (𝑝 → 𝑙𝑜𝑔 ( ^𝑝^ )), motivated by Eq. 3. The value estimations were clamped between 0.01 and 0.99 to prevent infinite values. The boundaries for clamping did not qualitatively change our results. To capture any variations in value estimations that deviate from the objective feature values, we also fit linear mixed effects models using the identities of the two features and their interaction as the independent variables.

All mixed effect modeling was performed using the lme4 and lmerTest packages (Bates et al., 2015; Kuznetsova et al., 2017). By default, models with maximal random effect structure were fit. If the model failed to converge, we incrementally simplified the model by first removing the correlation parameters between random effects, and then any random slope or random intercept that had small standard deviations if it prevented convergence and did not significantly improve model fit. Differences between parameters were tested using contrast tests. A t-test was used for testing contrasts in linear mixed-effects models, and a Wald Chi-square test was used for generalized linear mixed-effects models (Fox & Weisberg, 2019).

### Computational models

#### Object-based RL

The object-based model assumes that participants learn the value of each object directly using a Rescorla-Wagner type reinforcement learning (RL) model. Because we provided reward feedback for both the chosen and unchosen options, we created a model that updates the values of both of the chosen and the unchosen stimuli. Moreover, we used separate learning rates for rewarded or unrewarded trials. This leads to four learning rates and the following update rules:

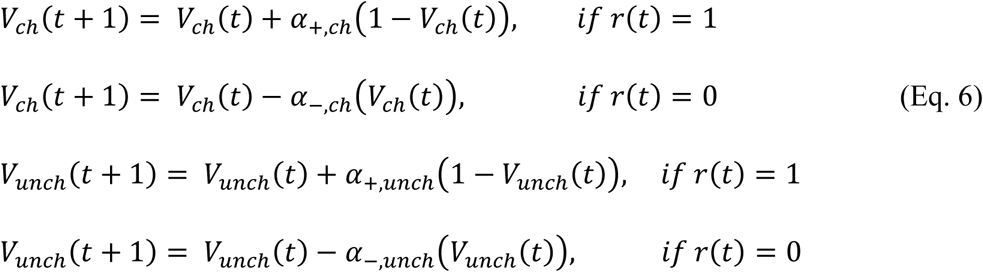

where 𝑉_*ch*_(𝑡) and 𝑉_*unch*_(𝑡) denote the values of chosen and unchosen options on trial *t*, *r*(*t*) is the reward outcome on trial *t* (1: rewarded, 0: unrewarded), *α*_+,*ch*_ and *α*_−,*ch*_ denote the learning rates for the chosen option on rewarded and unrewarded trials, and *α*_+,*unch*_ and *α*_−,*unch*_ denote the learning rates for the unchosen option on the rewarded and unrewarded trials.

We assumed that in each trial, the decision maker chooses between the two options based on the current estimate of reward probabilities of the two options. Hence, the probability of choosing option represented on the left (*Prleft*), over option represented on the right (*Prright*) was calculated as follows:

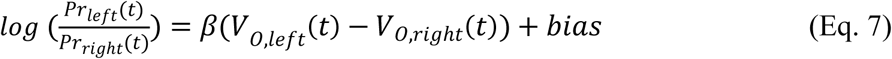

where 𝑉_𝑂,𝑙𝑒𝑓𝑡_(𝑡) and 𝑉_𝑂,𝑟𝑖𝑔ℎ𝑡_(𝑡) denote the values of the object on the left and right, 𝛽 is the inverse temperature with lower values (high temperature) resulting in more stochastic choice, and *bias* is the participant’s side bias in choosing between the two options.

It has been shown that the reward values of the unchosen object or feature(s) decay over time (Farashahi et al., 2017; Farashahi & Soltani, 2021; Niv et al., 2015). To capture this effect, we introduced a forgetting factor by adding a decay parameter to the RL models. More specifically. the reward probabilities (values) of the objects/features that were not updated due to being unavailable on a given trial decay to 0.5 with a constant rate of *d*:

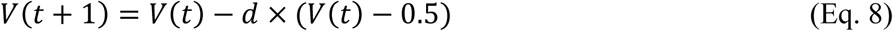

This model has 7 parameters in total, including one bias, an inverse temperature, four learning rates, and one decay parameters.

### Feature-based RL

The feature-based models learn the predictive values of each feature instance separately and then estimate the predictive value of each stimulus/object by combining these feature values. The update rule is similar to Eq. 6, applied separately to each feature, with potentially different learning rates for the two features (to capture attentional effects).

Subsequently, a weighted linear combination the feature values is used to predict choice as follows:

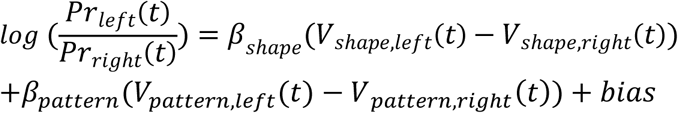

*Vshape* and *Vpattern* denote the values of feature shape and pattern. 𝛽_𝑠ℎ𝑎𝑝𝑒_ and 𝛽_𝑠ℎ𝑎𝑝𝑒_ denote the inverse temperatures for the shape and pattern values, which determine their influences on choice. Because sequence-prediction trials precede choice trials, we assumed that the effect of attention to be constant throughout the entire session. The full model, which includes separate learning rates and separate inverse temperatures for the two features, is referred to as the feature-based learning model with attention modulating both choice and learning (FtACL, 12 parameters). The models with only separate learnings rates or only separate inverse temperatures are referred to as feature-based learning model with attention modulating learning (FtAL, 11 parameters) and feature-based learning model with attention modulating choice (FtAC, 8 parameters), respectively. The base model uses the same learning rates and inverse temperatures for both features (denoted Ft, 7 parameters).

### Model fitting and model selection

We fit the above RL models (and their variants) to the participants’ choice behavior. The models were fit to the experimental data using a hierarchical Bayesian approach (Ahn et al., 2017) implemented in NumPyro (Bingham et al., 2019; Phan et al., 2019). The parameters were sampled using the No-U-Turns sampler, an efficient variant of the Markov chain Monte Carlo (MCMC) algorithm. Specifically, four chains were run in parallel and 3000 samples were taken from each chain. The first 1000 from each were used as warm-up samples and discarded from further inference, resulting in a total of 8000 posterior samples. The hierarchical Bayesian approach aggregates information from multiple participants to inform the group-level parameter estimates while also using the group-level parameter estimates to regularize estimates of individual parameters (Ahn et al., 2017), making it a good model fitting methodology for fitting complex cognitive models where each participant goes through a limited number of trials.

Following recommendations by Ahn et al. (2017), a group-level prior of Normal(0, 1) was put on the means of all the parameters except for the bias, which had a prior of Normal(0, 10). A group-level prior of HalfCauchy(0, 10) was put on the standard deviation of all the parameters. The participant-level parameters were reparametrized by shifting and scaling standard normal random variables with the group means and standard deviations. All participant-level learning rates and decay rates were then put through an inverse probit transform to be bound between 0 and 1, and all participant-level inverse temperatures were put through an exponential transform to ensure positivity. For the control analyses with tied learning rates (the FA model in **Fig. S3A**), the effect of attention was modeled as an additional parameter being added to the learning rates of the feature with regularity before the inverse probit transform. It had the same prior as all other learning rate parameters. All parameters had an r-hat less than or equal to 1.01, effective sample size of at least 400, and no model had divergence.

We compare the Watanabe-Akaike information criterion (WAIC) between different models as a measure of the goodness-of-fit and the stacking weights (Vehtari et al., 2017; Yao et al., 2018). WAIC approximates the expected out-of-sample predictive performance of a model using its in-sample performance and complexity, which includes the effective number of parameters (Vehtari et al., 2017). Stacking weights optimally combine different models to achieve the highest predictive performance (Yao et al., 2018). Models that are more effective at prediction are assigned higher stacking weights. WAIC was calculated using leave-one-subject-out approximate cross validation, except for trial-wise WAIC. We consider an improvement of WAIC bigger than 1.96 times the standard error to be significant (Vehtari et al., 2017). Model comparisons and parameter estimations were computed using the ArviZ package (Kumar et al., 2019). We used 1000 randomly selected posterior samples for model validation simulations, a method referred to as the posterior predictive check (Ahn et al., 2017; Baribault & Collins, 2023). We found that increasing the number of samples did not qualitatively change the results.

### Parameter estimation and hypothesis testing

To perform hypothesis testing on the value of a given parameter (Makowski et al., 2019), we report the median of the posterior samples of that parameter as a measure of central tendency. Furthermore, we report the probability of direction, which is the proportion of posterior samples that are positive or negative, whichever is higher. This is analogous to one minus the one-sided p-value. We interpret a probability of direction greater than 0.95 as evidence that a parameter is significantly greater than or smaller than 0. We also report the 95% highest posterior density interval (95% HPDI), which is the interval that contains 95% of the posterior samples, and such that points within the interval have a higher probability density than points outside of it.

All hypothesis testing on parameters were performed using the posterior distribution of the group-level mean parameters unless otherwise noted. We defined positivity and negativity biases as the difference between the learning rates from reward and no reward for either the chosen or unchosen options. We defined the effect of regularity manipulation (attention) as the difference between the learning rates for the same choice and reward outcome (for example, chosen and rewarded, unchosen and rewarded, etc.) but separately for different features. Calculations of these effects were performed for each sample from the posterior distribution to arrive at the posterior distribution of the differences in parameter values, then the metrics in the previous paragraph were calculated.

### Model and parameter recovery

To validate the results of model comparison and parameter estimations, we performed both model recovery and parameter recovery. We used the posterior median of group mean and standard deviation as the true group-level parameters and sampled from a normal distribution specified by them to obtain the participant-level parameters. We used these parameters to simulate choice sequences using the same stimulus and reward sequences presented to participants in our experiment (N=57). Note that this type of post-hoc recovery analysis might be conservative because it uses model parameters that were obtained from fitting on the same dataset, potentially leading to more similar behavior between models. The individual differences of parameters with small posterior standard deviation or close to the boundary may be hard to identify. We consider a parameter well-recovered if its 95% HPDI includes the true parameter value used for simulation (Baribault & Collins, 2023). Using PSIS-LOO instead of WAIC as the model comparison metric (Vehtari et al., 2017) or having a narrower prior of HalfCauchy(0, 5) on the group-level standard deviation parameters did not qualitatively change the results of model and parameter recovery.

## Authors contributions

AS conceived the idea for the experiment. AY, MCW, ET, and AS designed the experiment. MCW and ET developed the computer code to run the experiment. AY and MCW conducted the experiment. AY, MCW, and AS developed the computational models. AY and MCW implemented the computational models and performed model simulations. AY, MCW, and AS analyzed the experimental data and simulations. AY, MCW, and AS wrote the first draft of the manuscript. All authors edited the manuscript.

### Acknowledgements

We would like to thank Daniela Garcia for assistance with the experimental design, Elizabeth Frey for help with the experimental design and for developing the code to run the experiment, and Marissa Benz for conducting the experiment. This work was supported by a National Science Foundation CAREER Award (BCS1943767) to A.S.

## Supplementary materials

**Figure S1.**
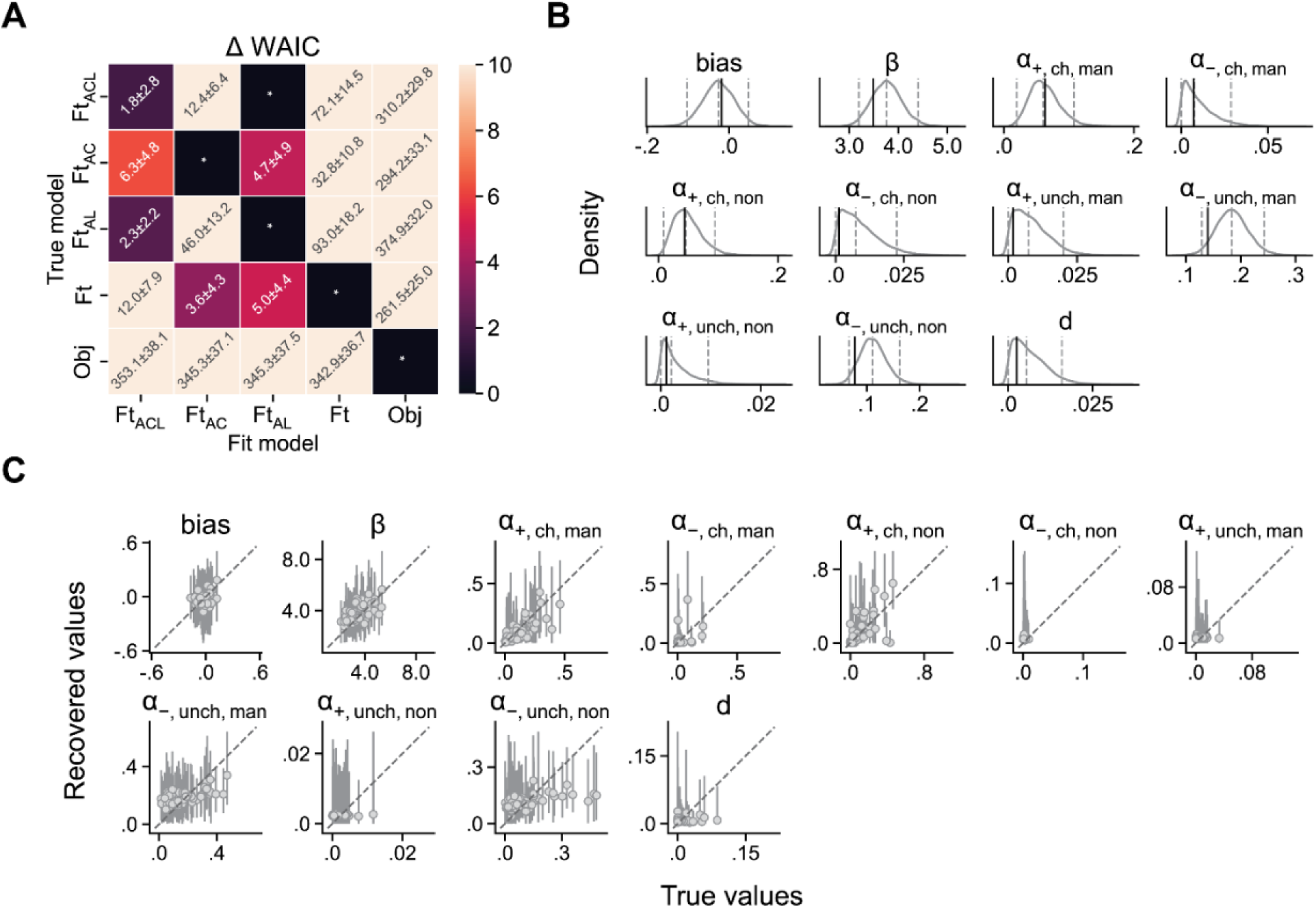
Model and parameter recovery. (**A**) Results of model recovery. Reported are the WAIC difference ± SE between each model and the best-fit model. An asterisk marks the best-fit model, which accurately identifies the correct model in all cases except for the Ft_ACL_ model. Although some feature-based models may be mistaken for each other, no incorrect model provided a significantly better fit than the true model. (**B**) Recovery of group-level mean parameters shows that the 95% HPDI of all parameters includes the true parameter values, indicating an acceptable level of recovery. (**C**) At the participant level, the 95% HPDI of 96.97% of the recovered parameters includes the true parameter values, confirming an overall acceptable level of recovery.

**Figure S2.**
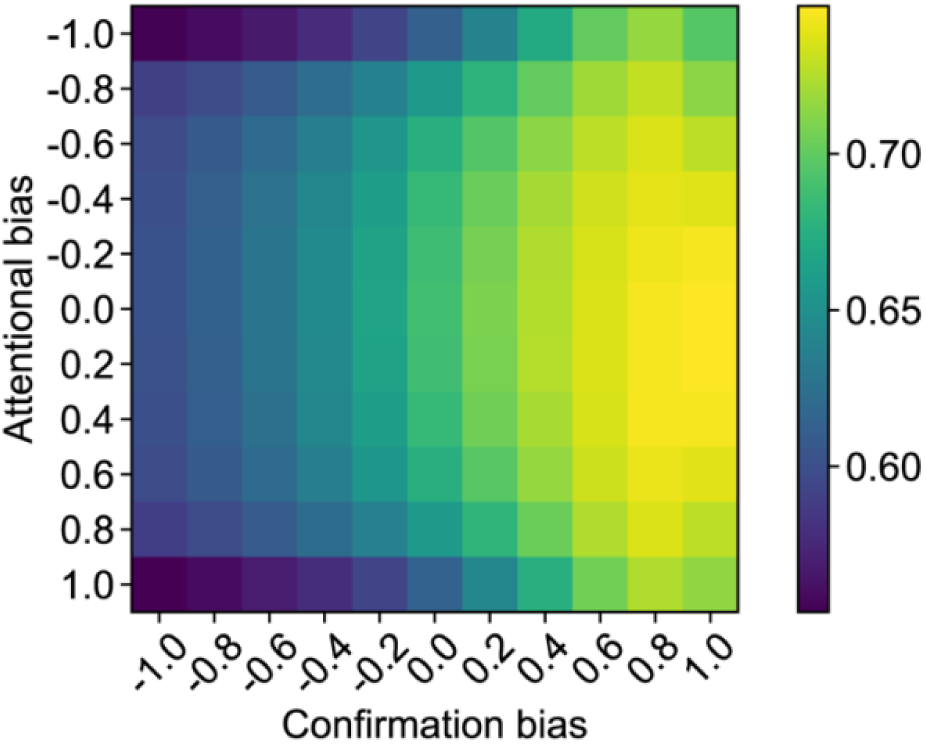
Performance of RL models with different levels of attentional and confirmation bias. The color in the heatmap shows the performance of a models simulated on the same 57 stimuli sequences shown to participants. All parameters were set to the median of the posterior group mean of the Ft_AL_ model, except for the learning rates. We set an average learning rate to be 0.1 for our simulations, which is close to the empirical average learning rate. We varied the levels of confirmation bias (*Conf*) and attentional bias (*Attn*), such that for one feature, *α*_*ch*,+_ = *α*_*unch*,−_ = 0.1(1 + 𝐴𝑡𝑡𝑛)(1 + 𝐶𝑜𝑛𝑓), and *α*_*ch*,−_ = *α*_*unch*,+_ = 0.1(1 + 𝐴𝑡𝑡𝑛)(1 − 𝐶𝑜𝑛𝑓), and for the other feature, *α*_*ch*,+_ = *α*_*unch*,−_ = 0.1(1 −𝐴𝑡𝑡𝑛)(1 + 𝐶𝑜𝑛𝑓), *α*_*ch*,−_ = *α*_*unch*,+_ = 0.1(1 − 𝐴𝑡𝑡𝑛)(1 − 𝐶𝑜𝑛𝑓). This parameterization allows us to examine the optimality of having these two types of biases in learning, both of which we empirically observe. We found that a strong confirmation bias leads to improved performance by enhancing learning from both chosen and rewarded options, as well as unchosen and unrewarded options, while ignoring other outcomes. In contrast, an attentional bias toward either feature results in decreased performance

**Figure S3.**
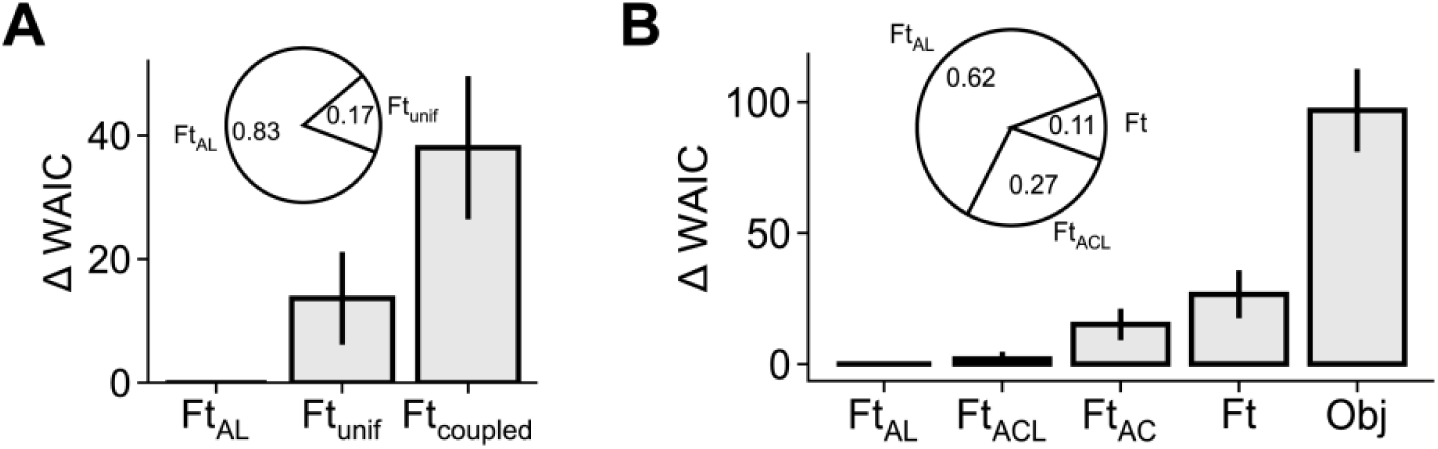
Supplementary model comparisons show that attention had distinct effects on learning from chosen and unchosen options, and the effect of feature-based attention was stronger in the first half of trials. (**A**) The plot compares of the goodness-of-fit between the full Ft_AL_ model (denoted “Full”) and two variations that lack some of the components, measured by the difference in WAIC between each model and the full model. The pie chart shows the stacking weights calculated using WAIC. Error bars show 1 standard error. The full model assumes separate learning rates for each feature, rewarded and unrewarded trials, and for chosen and unchosen options (total 8 learning rates). The feature-based model with uniform effect of attention (Ft_unif_) model simplifies this by treating the effect of attention as a single parameter that modulates all 4 learning rates uniformly. The feature-based model with coupled learning rates (Ft_coupled_) assumes separate learning rates for confirmatory or disconfirmatory outcomes and for each feature (total 4 learning rates). Both simplifications led to worse fit compared with the full model (*ΔWAIC* = 13.65 ± 7.50 and 38.01 ± 11.58). A weighted combination of the full Ft_AL_ model and the feature-attention (FA) model achieved the highest predictive performance, with the full Ft_AL_ model having larger weight (inset pie chart). (**B**) Model comparison based on fitting choice data from the first half of trials using different models. The plot compares the goodness-of-fit of different models, measured by the difference in WAIC between each model and the best-fitting model (FtAL). The pie chart shows the stacking weights calculated using WAIC. Conventions are the same as in Fig. 4. Error bars show 1 s.e.m. Feature-based learning models that include attention modulating learning have larger weights than models without this feature and moreover, participants did not utilize object-based learning in the first half of the experiment.

**Figure S4.**
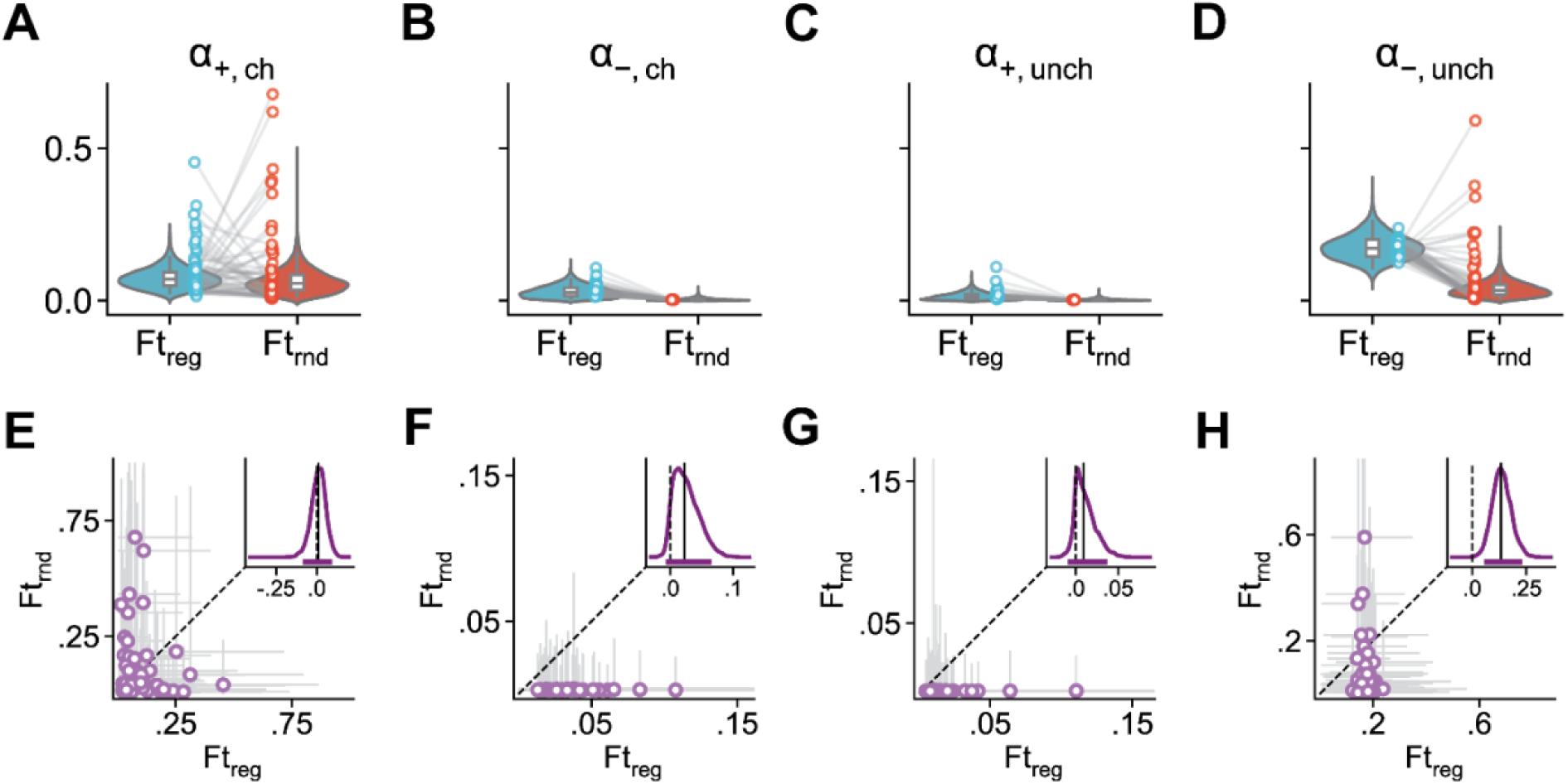
Attentional modulation enhanced learning the values of the feature with regularity in the first half of the experiment. (**A–D**) Evidence of strong confirmation bias and selective attention in feature-based learning. Violin plots show posterior distributions of group-level learning rates for chosen (ch) and unchosen (unch) options and separately for rewarded (*α*_+_) and unrewarded (*α*_−_) trials. Each point shows the learning rates for individual participant and whiskers of the box plot show the 95% HPDI. (**E–H**) Evidence of regularity manipulation influencing the learning rates, particularly about unchosen and unrewarded options, which was stronger than the effect when fitting the model on all the trials (compare with Fig. 5D, H). Scatter plots show the posterior distributions of individual participants’ learning rates. Dots show the median and grey bars show the 95% HPDI. The inset plot shows the posterior distribution of the difference between the group-level learning rates for Ft_reg_ and Ft_rnd_. Vertical solid line indicates the median. Dotted line indicates zero. The purple band shows the 95% HPDI.

**Figures S5.**
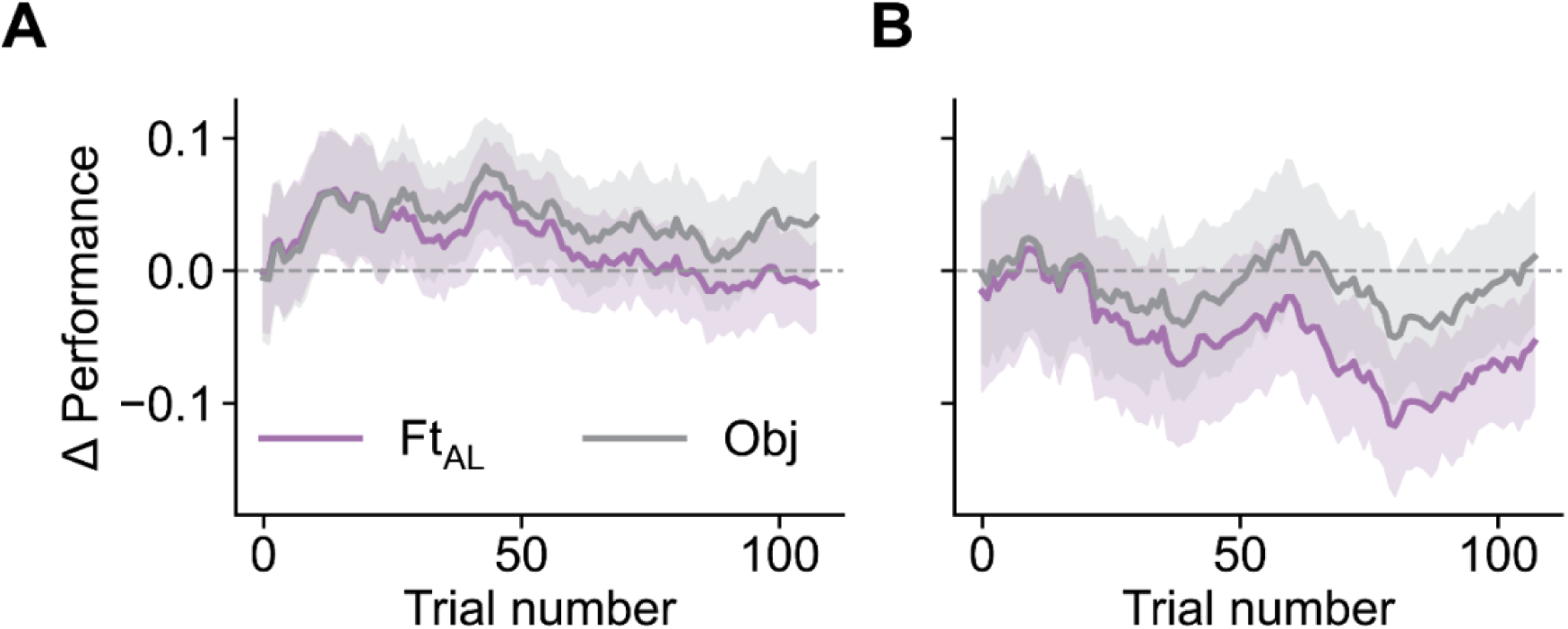
Differences between simulated and empirical learning curves, shown separately for participants whose choice behavior was better fit by the feature-based model (Ft_AL_, *N*=38) in panel (**A**), and the object-based model (Obj, *N*=19) in panel (**B**). Participants better described by the Ft_AL_ model were consistently outperformed by the Obj model, but their performances were more similar to the Ft_AL_ model. Participants better described by the Obj model outperform the Ft_AL_ model, but their performances were more accurately matched by the Obj model. Overall, this indicates that between-participant variability in learning strategies (feature-based vs. object-based) partially explains the results shown in Fig. 5C.

**Figure S6.**
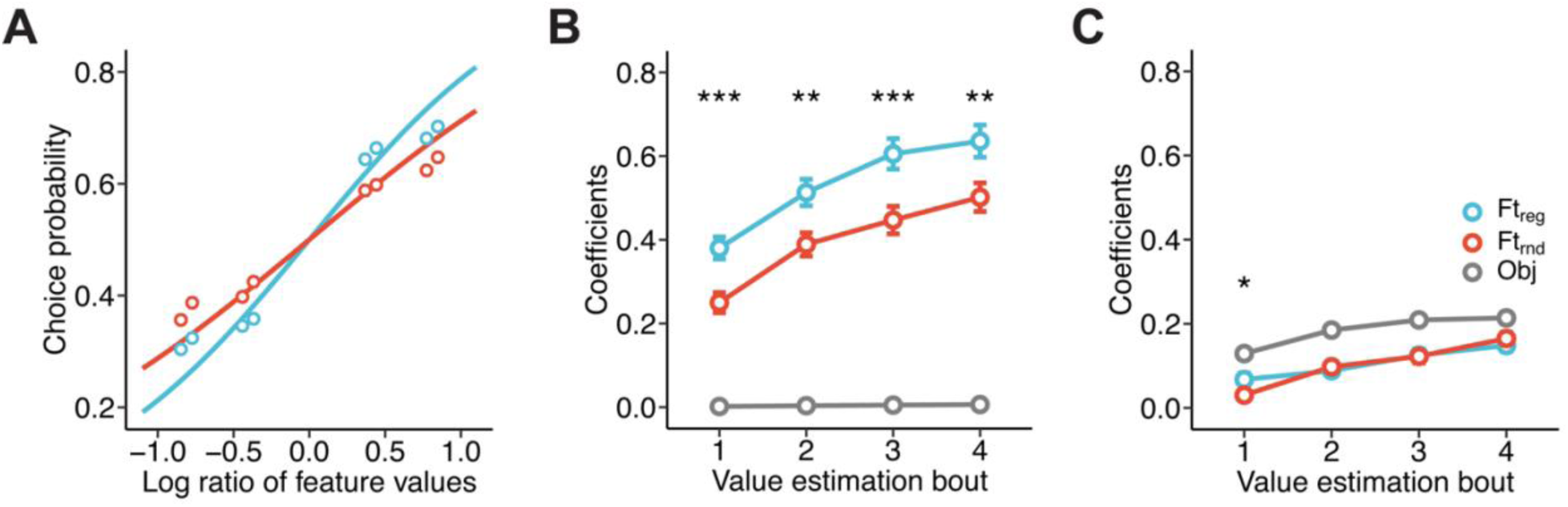
Posterior predictive checks of the RL models replicate empirical behavioral signature in choice and estimation. (**A**) Results of mixed-effects logistic regression using the log ratio of feature values to predict simulated choice. Each point represents the choice probability associated with a specific log feature value ratio. Error bars show one s.e.m., and the lines show the best fit logistic curves. The slope for the feature with regularity (Ft_reg_) was significantly higher than that of the random feature (Ft_rnd_) (*χ^2^*(1)=15.21, *p*=9.60×10^-5^). (**B-C**) Results of a mixed-effects linear regression using feature and object values (Obj) to predict RL values in four bouts of value estimation trials, separately for the Ft_AL_ model (**B**) and Obj model (**C**). Each point represents the fixed effects regression coefficient, with error bars indicating the standard error. Asterisks above the points indicate the significance of the contrast between the feature with regularity and random feature (\**p*<0.05; \*\**p*<0.01; \*\*\**p<0.001*). The Ft_AL_ model’s value estimations were a weighted sum of feature values, with a higher weight placed on Ft_reg_. The Obj model’s value estimations were more consistent with stimulus/object values.

**Figure S7.**
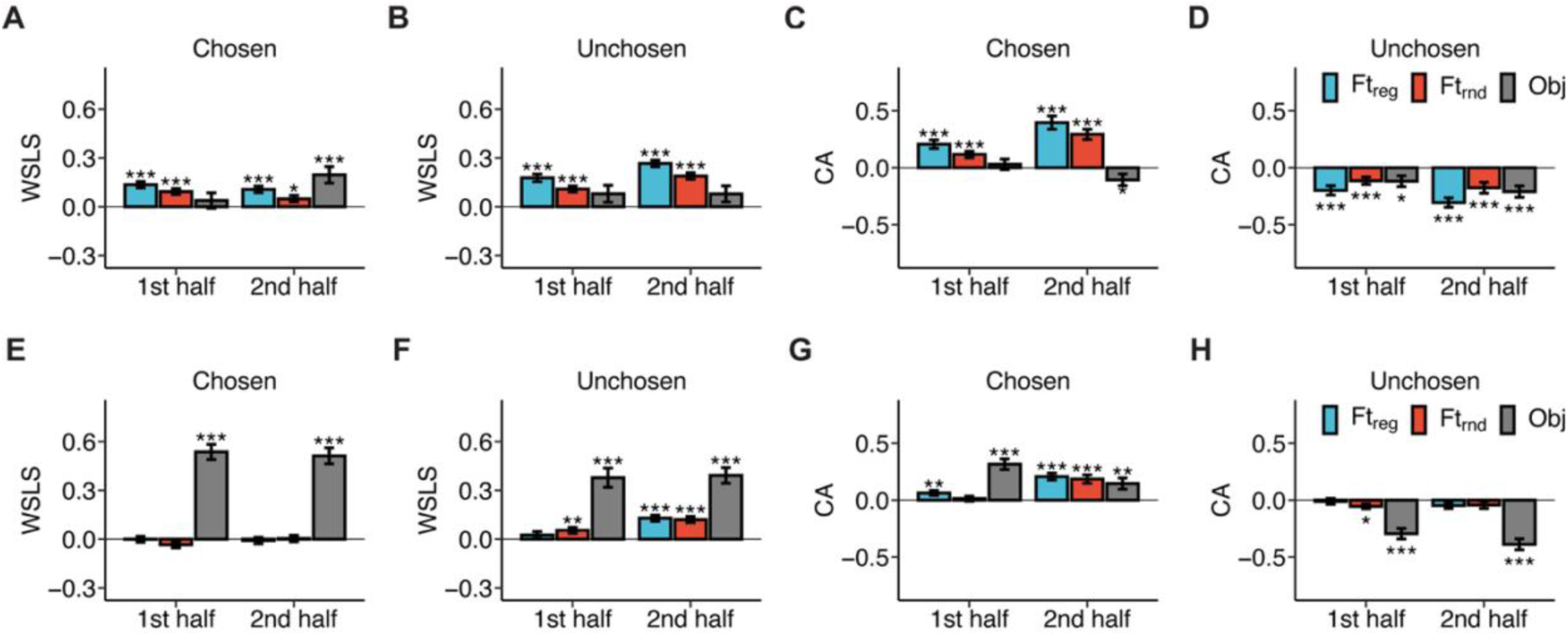
Posterior predictive checks of the RL models replicate behavioral signatures in WSLS and CA strategies. (**A–D**) WSLS and CA based on the chosen and unchosen options, calculated from the simulated behavior of the Ft_AL_ model. (**E–H**) WSLS and CA based on the chosen and unchosen options, calculated from the simulated behavior of the Obj model. Error bars represent 1 standard error, and asterisks indicate the significance level of the coefficient’s difference from zero (\**p* < 0.05, \*\**p* < 0.01, \*\*\**p* < 0.001). Ft_reg_, Ft_rnd_ and Obj denote the feature with regularity, random feature, and object, respectively.

**Table S1.**
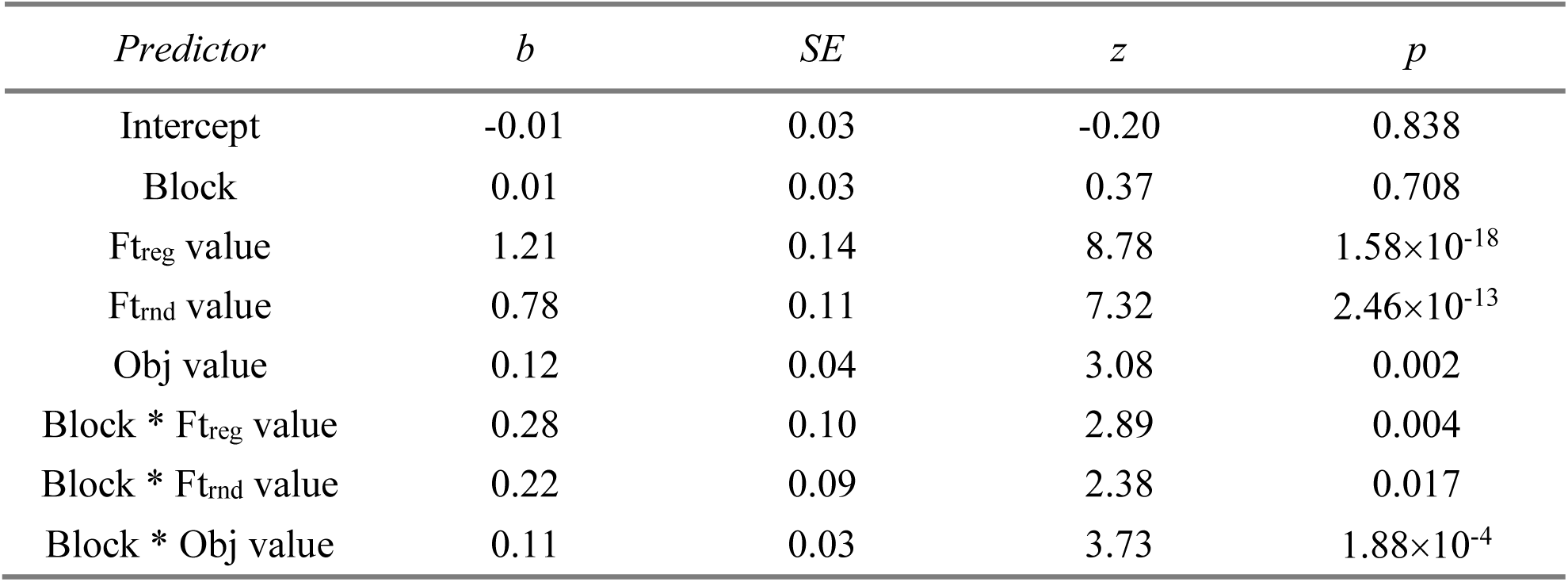
Results of fitting choice data with a generalized linear mixed-effects model based on log odds of feature and object values and block as independent fixed factors. Ft_reg_, Ft_rnd_ and Obj denote the feature with regularity, random feature, and object, respectively. The block factor denotes whether the trial was in the first or second half of the session. Interaction between block and the value predictors reflect changes in knowledge about feature and object values due to learning.

| *Predictor* | *b* | *SE* | *z* | *p* |
| --- | --- | --- | --- | --- |
| Intercept | -0.01 | 0.03 | -0.20 | 0.838 |
| Block | 0.01 | 0.03 | 0.37 | 0.708 |
| Ft<sub>reg</sub> value | 1.21 | 0.14 | 8.78 | 1.58×10<sup>-18</sup> |
| Ft<sub>rnd</sub> value | 0.78 | 0.11 | 7.32 | 2.46×10<sup>-13</sup> |
| Obj value | 0.12 | 0.04 | 3.08 | 0.002 |
| Block * Ft<sub>reg</sub> value | 0.28 | 0.10 | 2.89 | 0.004 |
| Block * Ft<sub>rnd</sub> value | 0.22 | 0.09 | 2.38 | 0.017 |
| Block * Obj value | 0.11 | 0.03 | 3.73 | 1.88×10<sup>-4</sup> |

**Table S2.** Results of fitting participants’ estimated values with a linear mixed-effects model based on log odds of feature and object values as independent fixed factors, separately for each bout of estimation trials. Ftreg, Ftrnd, and Obj denote the feature with regularity, random feature, and object, respectively. Ft_reg_-Ft_rnd_ denotes the result of the contrast test.

| *Bout* | *Predictor* | *b* | *SE* | *t* | *df* | *p* |
| --- | --- | --- | --- | --- | --- | --- |
| 1 | Intercept | 0.05 | 0.04 | 1.14 | 797.60 | 0.256 |
|  | Ft<sub>reg</sub> | 0.58 | 0.10 | 5.68 | 72.74 | 2.65×10<sup>-7</sup> |
|  | Ft<sub>rnd</sub> | 0.24 | 0.08 | 2.98 | 84.45 | 0.004 |
|  | Obj | -0.02 | 0.03 | -0.67 | 797.60 | 0.502 |
|  | Ft<sub>reg</sub>-Ft<sub>rnd</sub> | 0.33 | 0.12 | 2.76 | 108.99 | 0.007 |
| 2 | Intercept | 0.09 | 0.04 | 2.22 | 797.76 | 0.027 |
|  | Ft<sub>reg</sub> | 0.46 | 0.08 | 5.52 | 83.20 | 3.77×10<sup>-7</sup> |
|  | Ft<sub>rnd</sub> | 0.22 | 0.08 | 2.70 | 84.01 | 0.008 |
|  | Obj | 0.09 | 0.03 | 3.59 | 797.76 | 3.47×10<sup>-4</sup> |
|  | Ft<sub>reg</sub>-Ft<sub>rnd</sub> | 0.24 | 0.11 | 2.21 | 116.88 | 0.029 |
| 3 | Intercept | 0.12 | 0.04 | 3.15 | 797.36 | 0.002 |
|  | Ft<sub>reg</sub> | 0.47 | 0.09 | 5.50 | 80.68 | 4.29×10<sup>-7</sup> |
|  | Ft<sub>rnd</sub> | 0.25 | 0.07 | 3.53 | 98.19 | 6.26×10<sup>-4</sup> |
|  | Obj | 0.14 | 0.03 | 5.61 | 797.36 | 2.74×10<sup>-8</sup> |
|  | Ft<sub>reg</sub>-Ft<sub>rnd</sub> | 0.22 | 0.10 | 2.24 | 110.19 | 0.027 |
| 4 | Intercept | 0.12 | 0.03 | 3.45 | 797.92 | 5.83×10<sup>-4</sup> |
|  | Ft<sub>reg</sub> | 0.52 | 0.08 | 6.10 | 73.93 | 4.44×10<sup>-8</sup> |
|  | Ft<sub>rnd</sub> | 0.40 | 0.06 | 6.34 | 92.80 | 8.06×10<sup>-9</sup> |
|  | Obj | 0.12 | 0.02 | 5.58 | 797.92 | 3.30×10<sup>-8</sup> |
|  | Ft<sub>reg</sub>-Ft<sub>rnd</sub> | 0.11 | 0.10 | 1.17 | 104.97 | 0.245 |

**Table S3.**
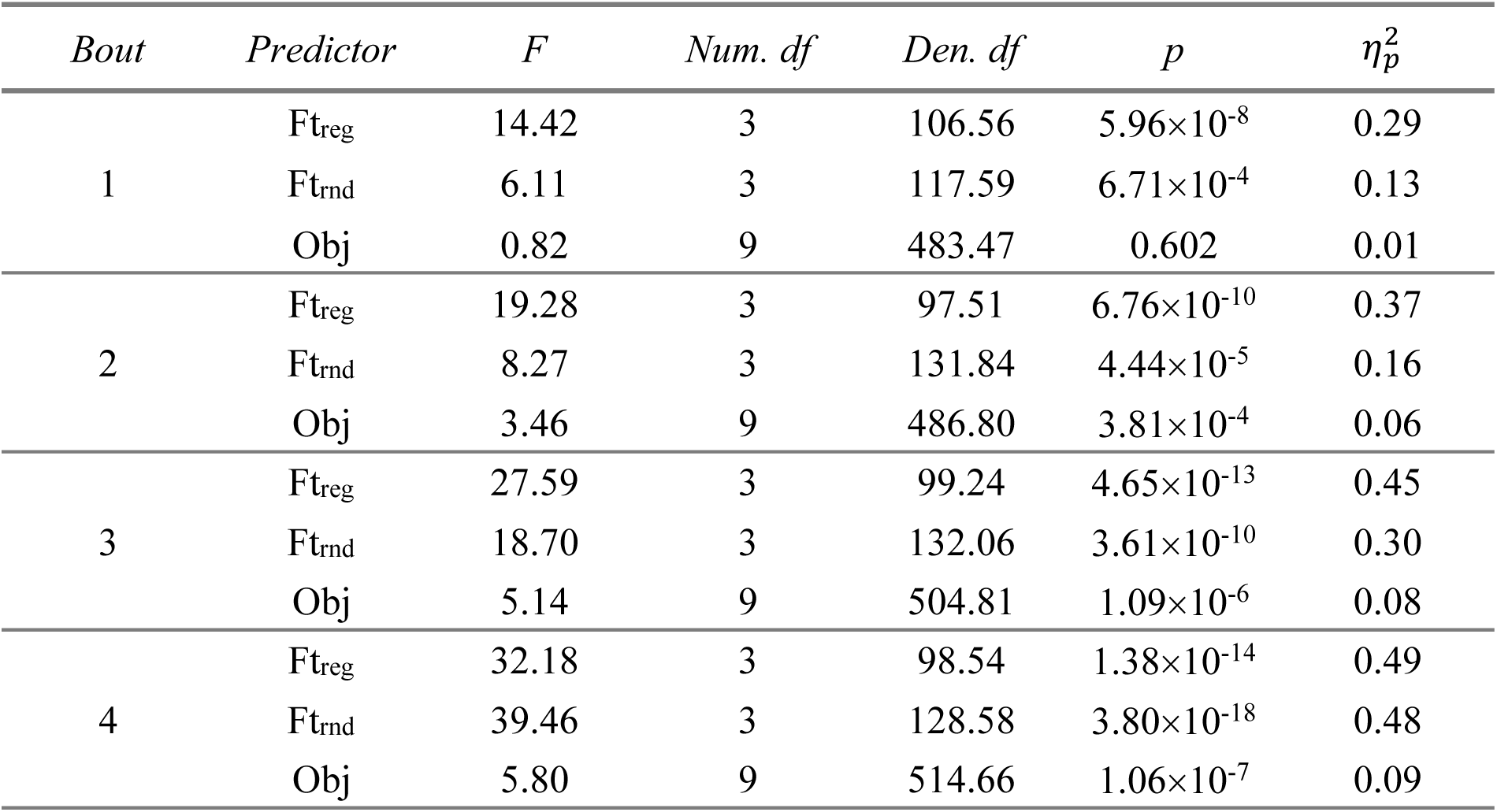
Results of fitting participants’ estimated values with a linear mixed-effects model based on feature and object identities as fixed factors, separately for each bout of estimation trials. Ft_reg_, Ft_rnd_, and Obj denote the feature with regularity, random feature, and object, respectively. The object effect is the same as interaction between the features because each stimulus can be identified as a conjunction of two features.

**Table S4.**
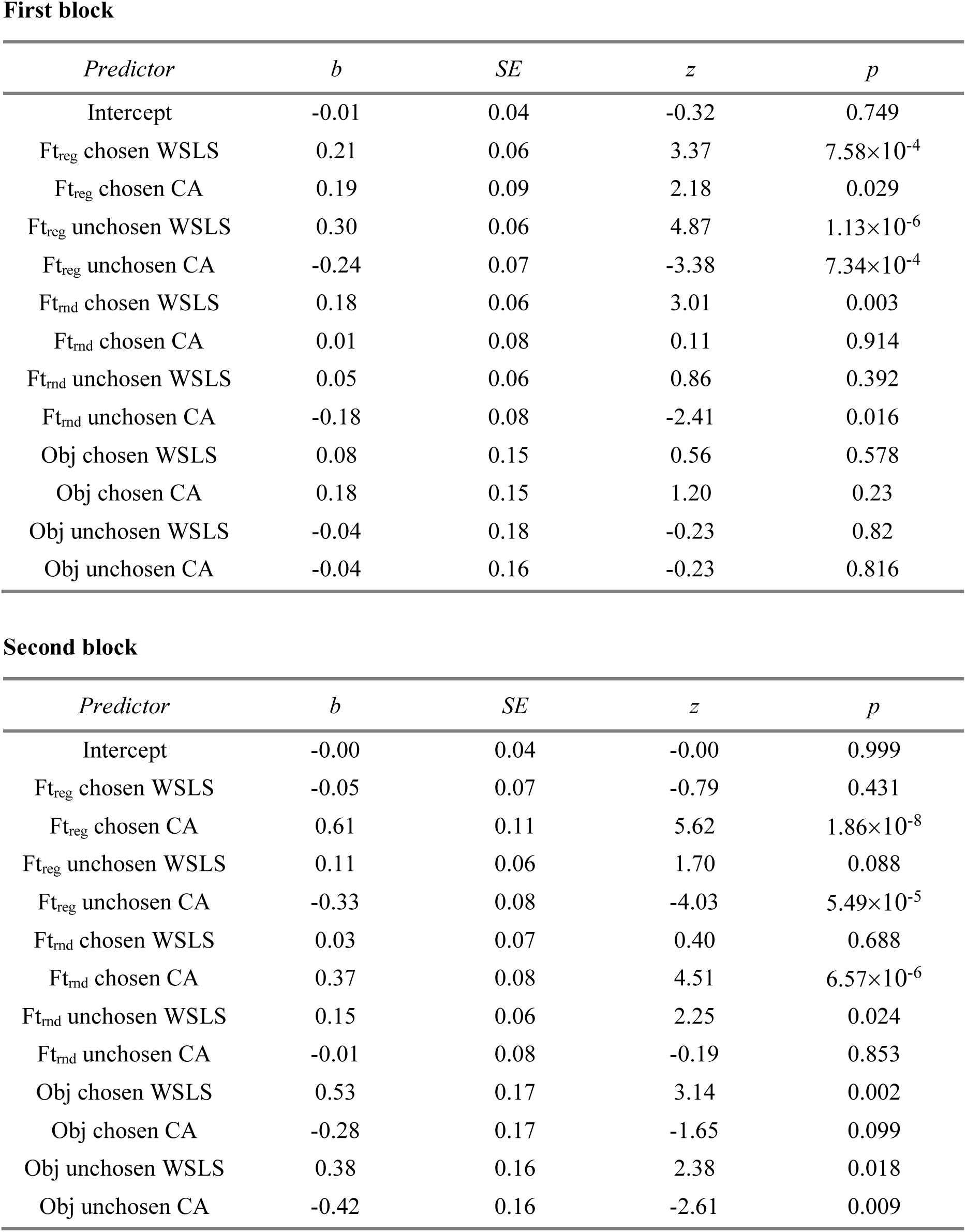
Result from generalized linear mixed-effects model predicting choice on a trial-by-trial basis based on reward and choice outcomes on the previous trial. The model incorporates individual features and stimulus identity, separately for the first and second block of the experiment. WSLS: win-stay-lose-switch, CA: choice autocorrelation.

**First block**
| *Predictor* | *b* | *SE* | *z* | *p* |
| --- | --- | --- | --- | --- |
| Intercept | -0.01 | 0.04 | -0.32 | 0.749 |
| Ft<sub>reg</sub> chosen WSLS | 0.21 | 0.06 | 3.37 | 7.58×10<sup>-4</sup> |
| Ft<sub>reg</sub> chosen CA | 0.19 | 0.09 | 2.18 | 0.029 |
| Ft<sub>reg</sub> unchosen WSLS | 0.30 | 0.06 | 4.87 | 1.13×10<sup>-6</sup> |
| Ft<sub>reg</sub> unchosen CA | -0.24 | 0.07 | -3.38 | 7.34×10<sup>-4</sup> |
| Ft<sub>rnd</sub> chosen WSLS | 0.18 | 0.06 | 3.01 | 0.003 |
| Ft<sub>rnd</sub> chosen CA | 0.01 | 0.08 | 0.11 | 0.914 |
| Ft<sub>rnd</sub> unchosen WSLS | 0.05 | 0.06 | 0.86 | 0.392 |
| Ft<sub>rnd</sub> unchosen CA | -0.18 | 0.08 | -2.41 | 0.016 |
| Obj chosen WSLS | 0.08 | 0.15 | 0.56 | 0.578 |
| Obj chosen CA | 0.18 | 0.15 | 1.20 | 0.23 |
| Obj unchosen WSLS | -0.04 | 0.18 | -0.23 | 0.82 |
| Obj unchosen CA | -0.04 | 0.16 | -0.23 | 0.816 |

**Table S4.** Result from generalized linear mixed-effects model predicting choice on a trial-by-trial basis based on reward and choice outcomes on the previous trial.
| *Predictor* | *b* | *SE* | *z* | *p* |
| --- | --- | --- | --- | --- |
| Intercept | -0.00 | 0.04 | -0.00 | 0.999 |
| Ft<sub>reg</sub> chosen WSLS | -0.05 | 0.07 | -0.79 | 0.431 |
| Ft<sub>reg</sub> chosen CA | 0.61 | 0.11 | 5.62 | 1.86×10<sup>-8</sup> |
| Ft<sub>reg</sub> unchosen WSLS | 0.11 | 0.06 | 1.70 | 0.088 |
| Ft<sub>reg</sub> unchosen CA | -0.33 | 0.08 | -4.03 | 5.49×10<sup>-5</sup> |
| Ft<sub>rnd</sub> chosen WSLS | 0.03 | 0.07 | 0.40 | 0.688 |
| Ft<sub>rnd</sub> chosen CA | 0.37 | 0.08 | 4.51 | 6.57×10<sup>-6</sup> |
| Ft<sub>rnd</sub> unchosen WSLS | 0.15 | 0.06 | 2.25 | 0.024 |
| Ft<sub>rnd</sub> unchosen CA | -0.01 | 0.08 | -0.19 | 0.853 |
| Obj chosen WSLS | 0.53 | 0.17 | 3.14 | 0.002 |
| Obj chosen CA | -0.28 | 0.17 | -1.65 | 0.099 |
| Obj unchosen WSLS | 0.38 | 0.16 | 2.38 | 0.018 |
| Obj unchosen CA | -0.42 | 0.16 | -2.61 | 0.009 |

**Table S5.** The influences of feature-based and object-based learning systems on participants’ value estimates. These influences are estimated by using a mixed-effects logistic regression model to predict participants’ value estimates based on reward probabilities from the feature-based with attention modulating learning (Ft_AL_) and object-based (Obj) models.

| *Bout* | *Predictor* | *b* | *SE* | *t* | *df* | *p* |
| --- | --- | --- | --- | --- | --- | --- |
| 1 | Intercept | 0.37 | 0.09 | 4.20 | 51.95 | 1.04×10<sup>-4</sup> |
|  | Ft<sub>AL</sub> | 1.23 | 0.13 | 9.15 | 49.34 | 3.30×10<sup>-12</sup> |
|  | Obj | 0.30 | 0.15 | 1.97 | 63.05 | 0.053 |
| 2 | Intercept | 0.29 | 0.07 | 3.98 | 78.67 | 1.53×10<sup>-4</sup> |
|  | Ft<sub>AL</sub> | 0.66 | 0.11 | 5.74 | 86.01 | 1.37×10<sup>-7</sup> |
|  | Obj | 0.63 | 0.13 | 4.97 | 63.09 | 5.47×10<sup>-6</sup> |
| 3 | Intercept | 0.26 | 0.05 | 4.85 | 89.98 | 5.18×10<sup>-6</sup> |
|  | Ft<sub>AL</sub> | 0.45 | 0.09 | 5.01 | 117.21 | 1.97×10<sup>-6</sup> |
|  | Obj | 0.77 | 0.09 | 8.22 | 705.90 | 9.39×10<sup>-16</sup> |
| 4 | Intercept | 0.31 | 0.06 | 5.06 | 75.50 | 2.90×10<sup>-6</sup> |
|  | Ft<sub>AL</sub> | 0.50 | 0.09 | 5.42 | 94.37 | 4.54×10<sup>-7</sup> |
|  | Obj | 0.71 | 0.08 | 8.37 | 63.63 | 7.52×10<sup>-12</sup> |

